# Untangling the effects of within-host and between-host natural selection in viruses with applications to evolution of HIV-1 co-receptor usage

**DOI:** 10.1101/2025.05.08.652803

**Authors:** Kieran O. Drake, Vinicius B. Franceschi, Fabrícia F. Nascimento, Erik M. Volz, the PANGEA Consortium

## Abstract

The fitness of a pathogen can be characterised at multiple levels corresponding to different stages of its life cycle, including replication within target cells (replicative capacity), immune evasion, and the capacity to infect new hosts (transmissibility). Pathogens evolve towards greater fitness within hosts, which can potentially compromise transmissibility. Typically, epidemiological studies for measuring pathogen fitness compute a single selection coefficient for pathogen variants that encompasses all stages of the pathogen life cycle. To address this problem, we develop a population-genetic model and software for the inference of multi-level selection coefficients which represent relative within-host fitness and transmissibility. Our approach builds on population genetic models previously applied to macro-evolution, whereby speciation and extinction rates are state-dependent. We relate the state-dependence in speciation and extinction rates to competing within- and between-host selection. Evolution of beneficial phenotypes within hosts may occur rapidly (higher ‘speciation’), but reduce rates of dispersal between hosts (higher ‘extinction’). This new coalescent-based method also accounts for highly nonlinear-population dynamics and variable sampling through time making it suitable for epidemiological studies. We derive the equilibrium proportion of a variant with multi-level selection under mutation-selection balance and show how this can be used in combination with the coalescent to greatly increase accuracy and precision of estimated selection coefficients. We characterise the accuracy of this method using a variety of birth-death and coalescent simulations before analysing a real dataset of HIV-1 genetic sequences from a large population-based sample of 4056 people living with HIV in Zambia. We quantified the multi-level relative fitness underlying HIV-1 ‘co-receptor switching’. In accordance with previous observations, we estimate that the relative transmissibility of X4-tropic virus is 9%, while within-host switching to X4-tropism is approximately seven-fold higher than switching to R5-tropism, indicating a strong within-host and between-host tradeoff of co-receptor usage. Our methods are available in an open-source R package musseco, providing a general-purpose tool for inferring multi-level selection coefficients for epidemiological sequence data.

**Author Summary:** The fitness of a pathogen can be characterised at each stage of its life cycle. However, epidemiological studies measuring pathogen fitness typically estimate a single value encompassing the entire life cycle. Pathogens evolve towards greater fitness within hosts, but this can compromise the capacity to infect new hosts. Our new method quantifies relative fitness at both levels. This method accounts for rapidly changing pathogen populations and variable sampling through time making it suitable for epidemiological studies. We tested our method using simulations and demonstrate its application by analysing real HIV-1 genetic sequences from people living with HIV in Zambia. This method sheds new light on long-standing questions about the relative fitness of HIV-1 variants which utilise different host-cell co-receptors. Using our new method, we estimate that the capacity of X4 variants to infect new people is just 9% of that of R5 variants, while within-host switching from R5 to X4 usage is seven times greater than the reverse switch. This indicates a strong within-host and between-host trade-off in co-receptor usage.

## 1 Introduction

Microbial evolution often features episodic natural selection that stems from competing selective pressures in fluctuating environments. The life cycle of a pathogen exhibits such a fluctuating environment, and requires the successful completion of multiple steps: Crossing tissue barriers, evading innate immunity, binding to a target cell, replicating within host cells, evading adaptive immunity, and shedding infectious particles such that the cycle may repeat. The selective pressures at each stage can conflict. For example, pathogen phenotypes optimised for one step (such as immune evasion) may perform poorly at others (like cell binding). Natural selection within hosts necessarily involves compromising some steps of the infection cycle to advance others.

Within-host and between-host evolutionary compromises are well-documented across diverse pathogens. The evolution of antimicrobial resistance often comes at the cost of reduced replicative fitness within host cells. Similarly, mutations enabling escape from adaptive immunity can impair receptor binding capacity [1]. A particularly illustrative example, which we examine in detail below, concerns HIV-1 co-receptor switching [2, 3, 4], such that different viral phenotypes experience varying selective advantages during the course of a single host infection. The use of the R5 co-receptor by viral variants to enter host cells is predominant at transmission and during all stages of infection. However, HIV-1 evolves during the course of infection to switch to the X4 co-receptor (or both co-receptors, i.e. dual tropic/R5×4) in up to 50% of individuals [2, 4], which is associated with a poorer baseline clinical status [5] and faster progression to AIDS [6, 7]. Although early studies suggested that R5 viruses were preferentially selected at transmission, as a result of a single or multiple leaky “gatekeepers” [8, 9], later transmission cluster analysis demonstrated that individuals infected by X4 viruses regularly transmit X4/R5×4, suggesting a stochastic process (random transmission hypothesis) [10, 11]. We address this question formally, and our findings support the existence of reduced transmissibility of X4 virus while indicating that limited transmission of X4 virus can still occur due to stochastic effects.

In quantitative terms, it is impossible to describe the fitness of a given phenotype using a single selection coefficient in the presence of multi-level selection. While multiple selection coefficients are needed to fully describe fitness variation across different environments, epidemiological studies have typically collapsed these into single estimates of variant fitness [12, 13]. And, despite the ubiquity of episodic selection in pathogens, there has been relatively little progress on methods to estimate multi-level selection coefficients of genomic variants. This is a challenging inference because of the large difference in timescales governing within-host pathogen replication and between-host transmission. Evolutionary rates within hosts are typically much higher than those measured between hosts, and the replication cycle of a pathogen between cells can be many orders of magnitude faster than the infection cycle between hosts. Another difficulty concerns the type of data available. Epidemiological surveillance data typically provides only one or a handful of sequences per host, but assessing the within-host fitness and transmissibility of particular variants would ideally make use of many sequences from hosts taken over the course of their infection which would itself be drawn from a population-based sample of cases. While deep sequencing data is enabling investigations into within-host microbial evolution [14, 15], such data are typically only collected at a single time point for each host.

In this paper, we present a general-purpose method for inferring multi-level selection coefficients for epidemiological sequence data. Our approach is similar to population genetic models previously applied in the macro-evolution setting, where multi-level pathogen selection parallels state-dependent speciation and extinction over evolutionary timescales [16]. In phylodynamic terms, competing within-host and between-host selection creates analogous state-dependence in speciation and extinction rates, but compressed into epidemiological timescales. Evolution of beneficial phenotypes within hosts may occur rapidly (higher ‘speciation’), but face reduced rates of dispersal between hosts (higher ‘extinction’). The key insight that enables inference of multi-level selection coefficients is that fitness depends on phenotypic state which changes over the phylogeny of the pathogen. Estimation of selection coefficients is thus coupled to reconstruction of ancestral phenotypes given the observed traits in sampled lineages.

Some limitations of previously-developed state-dependent selection models preclude their use for analysis of infectious diseases. On the short timescales of pathogen evolution, epidemiological dynamics need to be taken into account. Epidemic dynamics often leads to highly nonlinear changes in effective population size that cover many orders of magnitude. Furthermore, in contrast to studies of macro-evolutionary processes, sampling of microbes is highly distributed through time, and pathogen phylogenies often consist of a mixture of sampled lineages near the present and close to the most recent common ancestor. Sampling through time is often non-random, and the rate of sampling can vary drastically and unpredictably over time, since sampling is often non-systematic and responsive to emerging public health incidents.

In this study, we develop a coalescent-based method for inferring multi-level selection that accounts for highly nonlinear population dynamics and variable sampling through time. Our approach provides a general-purpose tool for inference of multi-level selection coefficients. It can be applied to any dataset where a time-scaled phylogeny can be estimated and where the phenotype (e.g. drug resistance, immune escape, or co-receptor usage) is known for each sample. This method is most appropriate for situations where one pathogen genome is sampled per host. It is optimised for analyses of pathogen genomes by automatically estimating effective population size over time (*N*_*e*_(*t*)), adjusting for nonlinear-population dynamics, allowing for heterochronous sampling, and conditioning on known sample dates.

## 2 Methods

### 2.1 The binary-state mutation and transmissibility model

In analogous fashion to the binary-state speciation and extinction (BiSSE) model [16], we specify the dynamics of two variants with different fitness at separate timescales. Our model is premised on the co-evolution of two variants 𝒜 and 𝒱 such that within hosts 𝒱 has a selective advantage, while also having lower transmissibility between hosts. The number of infected hosts with each variant over time is *Y*_*a*_(*t*) and *Y*_*v*_(*t*) and we will assume unless otherwise stated that the variants are in a mutation-selection balance, such that *Y* (*t*) = *Y*_*a*_(*t*) + *Y*_*v*_(*t*) and *Y*_*a*_(*t*) = *p*_*a*_*Y* (*t*) and *Y*_*v*_(*t*) = (1 − *p*_*a*_)*Y* (*t*) where *p*_*a*_ is the proportion which is of the ancestral type (c.f. section 2.2.2). The variants have different rates of transmission between hosts, with time-dependent per-capita rates *β*_*a*_(*t*) and *β*_*v*_(*t*). We parametrise the relative fitness of variant 𝒱 using

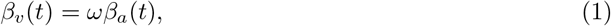

and assuming that a fitness penalty for 𝒱 such that *ω <* 1. We sometimes also refer to the *selection coefficient* for 𝒱 defined as *s* = *ω* − 1 and require that −1 *< s <* 0. We also assume that the generation time (mean time between successive infections) is the same between variants.

This model will require that each infected host at each time point is colonised by either 𝒜 or 𝒱 and that mutation from one variant to another is instantaneous. This model is therefore premised on within-host genetic diversity being negligible. We parametrised the increased within-host fitness of 𝒱 in terms of an imbalance between the rate of mutation from 𝒱 to 𝒜 and vice-versa. The mutation rate in a single host from 𝒜 to 𝒱 and from 𝒱 to 𝒜 are denoted *µ*_*av*_ and *µ*, respectively.

Our model is premised on increased mutation from 𝒜 to 𝒱 by a factor *α*:

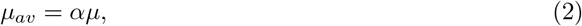

such that *α >* 1.

Describing multi-level selection therefore requires estimation of two parameters, *ω* and *α*. A compartmental diagram summarising the co-evolution of 𝒜 and 𝒱 is shown in Fig 1. In sections 2.2.2 and 2.2.1 we show how these parameters determined a mutation-selection balance between variants and how to account for epidemiological dynamics by appropriate choice of functional forms for *Y* (*t*) and *β*_*x*_(*t*). Figure 1 also illustrates genealogies simulated under different scenarios of strong and weak within-host and between-host selection. In the weak-selection scenario, clades of 𝒱 have longer persistence times and larger size, but under the constraint of a given mutation-selection balance, such a variant has correspondingly weak within-host selection leading to infrequent denovo occurrences within hosts. In contrast, under strong within- and between-host selection, 𝒱 occurs frequently throughout a genealogy, but clades of 𝒱 rapidly go extinct.

**Figure 1.**
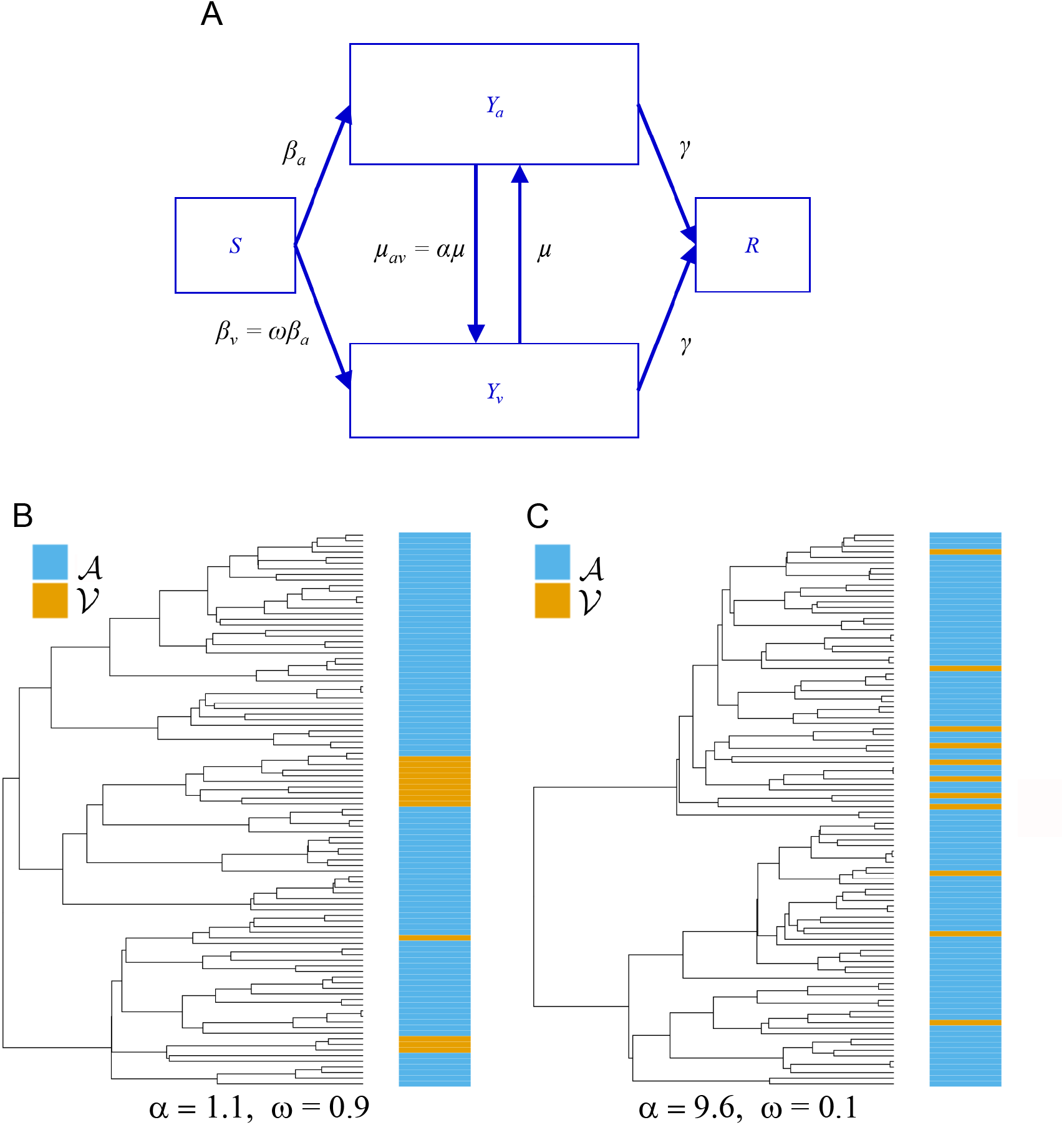
Binary-state mutation and transmissibility model. (A) Illustrative epidemiological compartmental model considering multi-level fitness. The model includes two variants, 𝒜 and 𝒱, which have different between-host transmission rates, with ratio *ω*, and asymmetric within-host mutation rates with ratio *α*. Panels (B) and (C) illustrate genealogies with identical mutation-selection balance of 𝒱 (≈ 15%), yet with different cladistic patterns stemming from different within- and between-host selection. Phylogenies with 100 tips were simulated with *diversitree* [17] and plotted using *ggtree* R package [18]. (B) Weak within- and between-host selection. The relative transmissibility of 𝒱 is 90% (*ω* = 0.9) and *α* = 1.1. (C) Strong within- and between-host selection. The relative transmissibility of 𝒱 is 10% (*ω* = 0.1) and *α* = 9.6.

### 2.2 The structured coalescent and modelling variant co-evolution

To model gene genealogies generated by the binary-state co-evolution model, we use the structured coalescent framework previously developed in [19] and [20]. This coalescent model specifies the rates of lineage migration and coalescence in terms of the corresponding population dynamics comprising three deterministic matrix-valued functions of time and the state of the system with *m* demes:

- Births: *F*_1:*m*,1:*m*_(*t, Y*). In this study, this represents between-host transmission rates for different pathogen variants.
- Mutation: *G*_1:*m*,1:*m*_(*t, Y*). In this study, this represents within-host mutation rates.
- Deaths: Γ_1:*m*_(*t, Y*). These terms may also correspond to recovery or removal from the infectious population in epidemiological models.

We order *m* = 2 demes corresponding to the co-evolving variants: D = (𝒜, 𝒱). Then, the *birth* matrix represents dispersal rates to new hosts and depends on the relative transmissibility parameter *ω*.

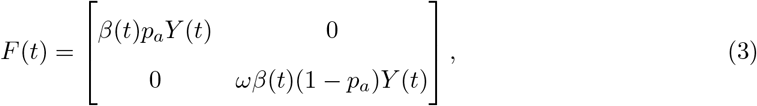

where *β*(*t*) is the time-varying per-capita transmission rate and *Y* (*t*) is the total number of infected hosts. We show below how *β*(*t*) and *Y* (*t*) are derived in terms of the effective population size over time. These birth rates appear on the diagonal since a host colonised with a particular variant may only generate new infections of the same type. In contrast, mutation terms appear on the off-diagonal and generate the simultaneous loss of a lineage of one type (𝒜 or 𝒱) and the gain of the other type:

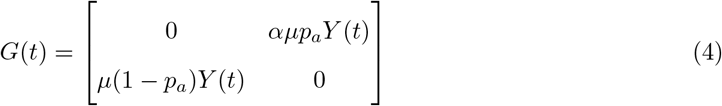

Finally, death rates determine the generation time of the process and allow translation of genealogical time to calendar time:

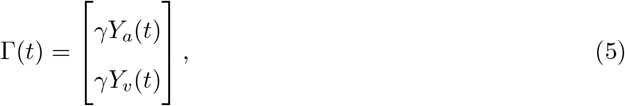

where the generation time *T*_*g*_ = 1*/γ* is assumed constant between variants.

In the supporting information, we review how the likelihood of a tip-labeled genealogy is computed conditional on the epidemiological history specified by ℳ = (*F* (*t, Y*), *G*(*t, Y*), Γ(*t, Y*)). With ℳ defined, the population size of each deme *k* can be computed by solving the system of ordinary differential equations:

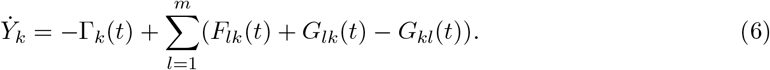

#### 2.2.1 Effective population size

To adjust for variation in the coalescent rate that is unrelated to the coevolution of 𝒜 and 𝒱, we calibrate *β*(*t*) and *Y* (*t*) so that the total coalescent rate (averaged over all demes) matches an empirical estimate that is obtained separately. Specifically, suppose that we are provided with an estimate of *N*_*e*_(*t*) for the entire gene genealogy that neglects population structure in the form the co-circulating variants. Our software implementation will estimate *N*_*e*_(*t*) using a non-parametric maximum likelihood method by default [21] (c.f. section 2.5), but in principle, any method can be used and provided as an input to our algorithm. Note that the scale of *Y* (*t*) is arbitrary, because the structured coalescent is over-determined, and many different forms of *β*(*t*) and *Y* (*t*) would yield the same *N*_*e*_(*t*). We need only to derive a definition such that changes in *Y* (*t*) yield identical changes in the pre-estimated *N*_*e*_(*t*). Our approximation to *Y* (*t*) is premised on the following:

1. Without loss of generality, *β*_*a*_(*t*) = 1, and transmissions *ϕ*_*a*_(*t*) per unit time from 𝒜 will be equal to population size *Y*_*a*_(*t*). For 𝒱, we will use *β*_*v*_(*t*) = *ω* and transmissions *ϕ*_*v*_(*t*) will be: *ϕ*_*v*_(*t*) = *ωY*_*v*_(*t*)
2. The coalescent rate for a pair of lineages in a deme *x* is based on the coalescent model in [19]: *λ*_*x*_(*t*) = 2*ϕ*(*t*)*/Y*_*x*_(*t*)^2^.

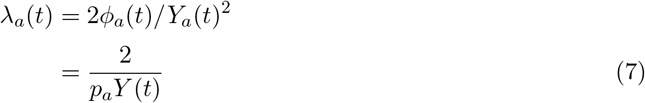

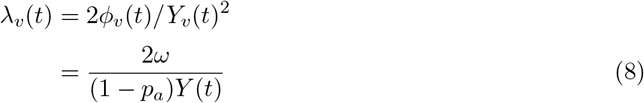
3. And, given the binomial probabilities 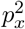 of selecting two lineages from the same deme *x*, the approximate pairwise coalescent rate for the population as a whole will be

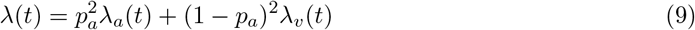

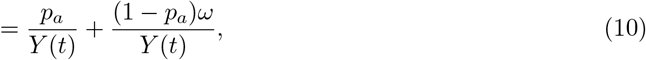

where the last line is obtained by substituting expressions from 7 and 8.

To solve for *Y* (*t*), we equate *λ*(*t*) (equation 10) to *ι/N*_*e*_(*t*), where *ι* is an additional scaling parameter which is estimated to improve the overall estimate of the coalescent rate. This yields

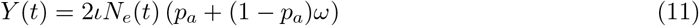

In practice, we observe that estimates of *ι* are very close to one (c.f. section 3.1), indicating that equation 11 is a good approximation in many situations.

#### 2.2.2 Mutation-selection balance

If the variant and ancestral types are in equilibrium (mutation-selection balance) it is possible to construct a likelihood of sampling proportions that are of the ancestral type (*p*_*a*_) and variant type (*p*_*v*_ = 1 − *p*_*a*_) assuming random sampling. This likelihood can be used in combination with the structured coalescent likelihood to gain additional precision when estimating selection coefficients, although the conditions of random sampling and mutation-selection balance should be evaluated first.

Here we solve for *p*_*a*_ by solving a set of detailed balance equations involving transmission, death (or recovery) and mutation. The corresponding rates are:

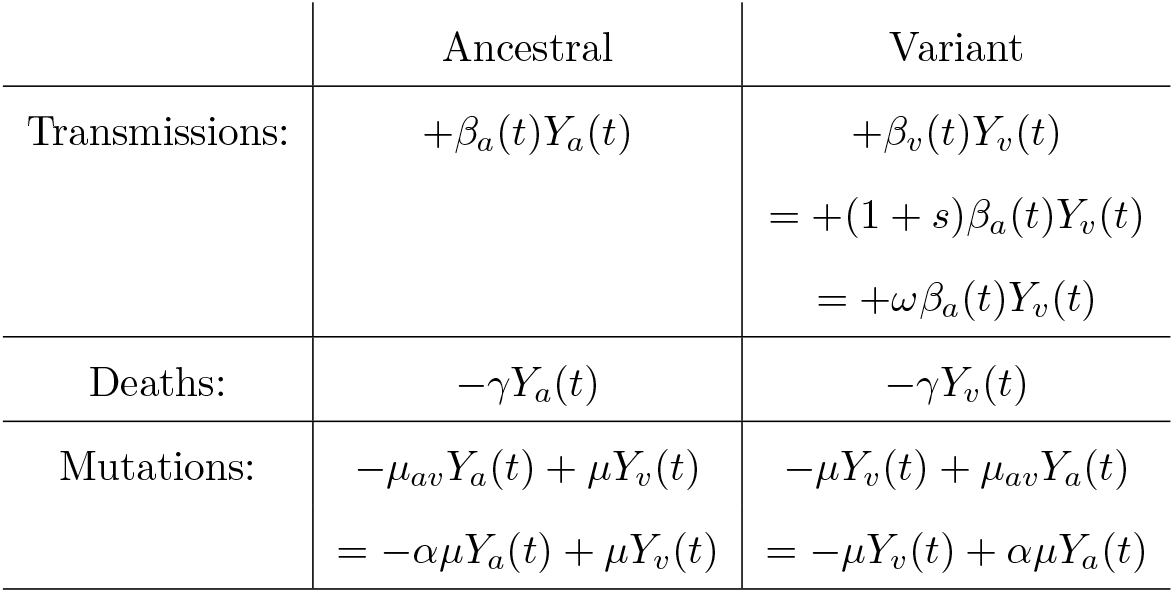

And, the following equations must be satisfied at equilibrium after dividing all expressions by *Y* (*t*) = *Y*_*a*_(*t*) + *Y*_*v*_(*t*):

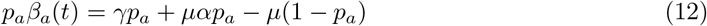

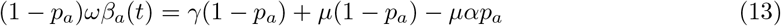

Combining these equations and solving for *p*_*a*_ yields

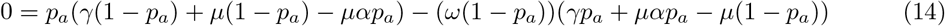

This quadratic equation in *p*_*a*_ has at most one root in the unit interval given the positive domain of all other parameters.

### 2.3 Simulations

We used three methods which support genealogical simulations in structured populations, *Coalescent*.*jl* [22], *TiPS* [23] and *diversitree* [17] to simulate trees in which we know the true values of the relative between-host fitness (*ω*) and relative within-host fitness (*α*) to test the new coalescent model. Two of these simulators, the *Coalescent*.*jl* [22] and *TiPS* [23], are based on the structured coalescent and require an epidemiological model describing the dynamics of infectious individuals. The *diversitree* method is based on a birth-death model and has previously been used to study the multi-state speciation and extinction process.

#### 2.3.1 Epidemiological model

To carry out simulations with *Coalescent*.*jl* and *TiPS*, we developed a mathematical epidemiological model to describe the dynamics of ancestral and variant states. This model was defined as two ordinary differential equations (ODEs) describing the number of infectious individuals carrying sequences with the ancestral state *Y*_*a*_(*t*) and the number of infectious individuals carrying sequences with the variant state *Y*_*v*_(*t*):

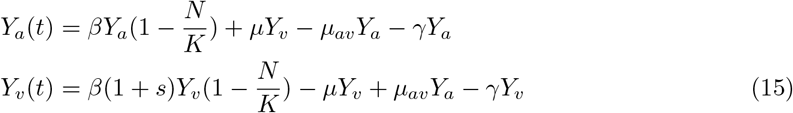

The rate of new infections was defined as logistic growth controlled by the parameter *β* for *Y*_*a*_(*t*) and *β*(1 + *s*), where *s* is the selection coefficient, for *Y*_*v*_(*t*). The carrying capacity was defined by the parameter *K* and was fixed at 10,000 individuals. Infected individuals recover at the rate *γ*.

Individuals containing the ancestral state, *Y*_*a*_(*t*), mutate to the variant state, *Y*_*v*_(*t*), at a rate *µ*_*av*_, and individuals carrying the variant state could revert to the ancestral state at a rate *µ*.

#### 2.3.2 Genealogical simulations

The parameter values used to simulate genealogies with *Coalescent*.*jl* (v0.1.4) and *TiPS* (v1.2.3) are described in Table S1. Both methods employ Gillespie’s direct algorithm with time-varying coalescent and migration rates. We obtained the value of the transition rate from *Y*_*a*_ to *Y*_*v*_ (*µ*_*av*_) using the theoretical equation for the mutation-selection balance using fixed values for the proportion of individuals in the ancestral state, the molecular clock rate (here also defined as the transition rate from *Y*_*v*_ to *Y*_*a*_), and the relative transmission fitness (*ω* = 1 + *s*).

To simulate trees with *Coalescent*.*jl* and *TiPS*, we also provided a sampling date randomly sampled in the past 5 years using the Julia programming language. We then used the same sampling date from *Coalescent*.*jl* trees to simulate trees using the *TiPS* R package.

For *diversitree* (v0.10.1) simulations, we analysed different scenarios of population dynamics with a high and low *R*_0_ (basic reproduction number). Note that *R*_0_ depends on the proportion of ancestral state and the birth rates. Each combination of parameter values analysed with *diversitree* had a different birth rate, but we grouped the results into two different values of *R*_0_: A high value of approximately 1.85 and a low value of approximately 1.22 (See Table S2 for a complete list of parameter values and the *R*_0_ for each combination of parameter values used with *diversitree*).

For all combinations of parameter values tested, we also sampled a total of 500 or 1000 individuals to understand how the sample size would influence the estimate of parameters.

#### 2.3.3 Simulation and analysis of genetic sequences

We carried out an additional set of simulations to mimic the observed HIV data described in section 2.4. We used *Coalescent*.*jl* with the same epidemiological model as described in section 2.3.1 to simulate genealogies in which trajectories were simulated for 29 or 45 years. For a list of the parameter values used, see Table S3. We simulated genealogies with 4056 tips with sampling time from the past 4 years (which was observed for the PANGEA dataset, see section 2.4). These simulated genealogies will be referred to as “true trees”.

We simulated sequence alignments of 105 bp and 2371 bp, based on the observed alignment length for the empirical dataset, using the simulated true trees and the program *AliSim* [24] to understand the impact of sequence alignment length on the estimation of relative fitness *α* and *ω*.

We simulated sequence alignments without gaps or recombination that started from random starting sequences. We then estimated maximum likelihood (ML) trees using the simulated sequence alignments and the program *IQ-TREE* (v2.4.0) [25], followed by the estimation of time-scaled trees using the strict clock implemented in *treedater* (v.0.5.3) R package.

#### 2.3.4 Statistical inference

Fitness parameters *ω* and *α* were estimated by maximum likelihood using prior knowledge of the generation time (equivalent to 1*/γ*) and the molecular clock rate of evolution, which we assume is equivalent to the reversion rate *µ*. Inference was carried out using both the coalescent and the augmented-coalescent likelihoods (see Supplementary Information for details). Approximate confidence intervals were computed assuming asymptotic normality of the maximum likelihood estimate (MLE) and computing standard errors from the observed Fisher information (negative Hessian of the log likelihood).

Performance of the estimator is summarised by relative error 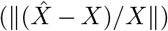 and the coverage probability of the 95% confidence interval.

### 2.4 Multi-level evolution of HIV-1 co-receptor tropism

In order to disentangle the effects of within- and between-host selection related to HIV-1 co-receptor usage, we used data from a high-prevalence setting in Sub-Saharan Africa. The HPTN 071 study, commonly referred to as PopART (Population Effects of Antiretroviral Therapy to Reduce HIV Transmission) was a large-scale community-based trial conducted in South Africa and Zambia between 2013 and 2018. Its primary aim was to assess the effect of a universal test-and-treat strategy on population-level HIV incidence. During the study period, ≈5% (n*>*48,000) of the total population of *>*1 million people were surveyed, including blood samples, annually for three years [26]. The HPTN 071-2 PopART Phylogenetics Ancillary study used samples from HIV seroconverters within the PopART cohort from Zambia only and HIV-positive samples collected from health clinics across nine of the 12 Zambian PopART communities, which were then subjected to sequencing and genome assembly processing as previously described [27]. Access to HIV-1 genomes and appropriate metadata was granted by the PANGEA-HIV 2 (Phylogenetics And Networks for Generalised Epidemics in Africa) Steering Committee after the acceptance of a concept sheet proposal.

#### 2.4.1 Data curation and phylogenetic tree reconstruction

Subtyping of all consensus sequences from this cohort was performed using *COMET* v2.4 [28]. Only the first sequence per individual and subtype C sequences (n=5460, 85.9%) were retained. The vast majority (85.3%) of those individuals were ART-naïve, and sequences from ART-experienced individuals were removed. Whole genome sequences (WGS) were aligned against the subtype B HXB2 reference using *MAFFT* v7.505 (auto and add-fragments options) [29]. Potential recombinants (*<*95% subtype C as determined by the Los Alamos National Laboratory [LANL] Recombinant Identification Program [*RIP*]) and hypermutated sequences were removed using the LANL Quality Control tool [30]. Finally, to ensure all data had a sufficient number of phylogenetically-informative sites, sequences shorter than 6500 ungapped nucleotides were removed, leading to n=4249 sequences.

Subsequently, HIV-1 genes were extracted. For the purposes of this work, which aimed to estimate multi-level selection of HIV-1 co-receptor usage, we retained *env* and V3 loop alignment subsets (alignment sizes of 2371 and 126 columns, respectively). For the complete *env* analysis, the highly variable loops V1 to V5 of *env* were masked following previous recommendations [31]. As simulations demonstrated (see Section 2.3.3), an alignment of approximately 100 nucleotides did not contain sufficient information to allow a reliable inference of *α* and *ω* parameters. Therefore, we do not report such fitness estimates derived from V3 sequences.

After testing multiple evolutionary models using *ModelFinder* v2.2.2.6 [32], we built an initial maximum likelihood (ML) tree using *IQ-TREE* v2.2.2.6 [25]. This analysis employed the general time reversible (GTR) model [33] with a free-rate model for rate heterogeneity across four categories (R4), iterating 100 times. A few sequences (n=7 for *env*) with *>*50% gaps or erroneous long branches in the ML tree were removed. After these curation steps, the *env* ML tree was re-built using: (i) the GTR nucleotide substitution model with empirical state frequencies (F), a proportion of invariable sites (I), and ten free-rate categories (R10); (ii) 1000 ultrafast bootstrap replicates; (iii) a perturbation strength of 0.2; and (iv) 3000 minimum iterations. Time-scaled phylogenetic trees were estimated after removing the HXB2 reference using an additive uncorrelated model [34] as implemented in *treedater* v0.5.3 [35] with 10 maximum iterations. A sub-sampled version of the time-scaled phylogenetic tree is shown in Fig S13 as an example.

#### 2.4.2 Co-receptor tropism and coalescent analyses

Nucleotide sequences spanning the V3 loop located in the gp120 genomic region inside *env* were extracted and submitted for co-receptor prediction using the *WebPSSM* server and a position- and subtype-C-specific SINSI scoring matrix [36, 37]. Subsequently, the distribution of *WebPSSM* scores, trends of co-receptor usage over time, distribution of CD4 counts and ranges for each predicted co-receptor, and the ratio of predicted R5 versus X4/R5×4 were computed.

For the coalescent analysis, the previously estimated *env* time-scaled tree was used alongside the co-receptor tropism predictions derived from the V3 loop region. A total of 4056 tips were considered, from which sample collection dates ranged from February 2014 to June 2018. The `fitbisseco` function from the *musseco* (v0.1.0) R package [38] was used. X4/R5×4 predicted sequences were treated as the variant state and R5 as the ancestral. The generation time (*T*_*g*_) was defined as 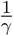, using the same *γ* value as in the simulations (Table S1). The molecular clock rate of evolution was based on the *treedater* estimate for *env* (3.2 × 10^*−*3^ nucleotide substitutions per site per year). The effective population size was internally estimated using the *mlesky* (v0.1.7) R package [21]. During this process, the smoothing parameter (*τ*) was optimised using cross-validation and uncertainty was quantified using parametric bootstrap (`parboot` function) with 100 replicates. The default initial values of the *α, ω* and *Y* (*t*) scale parameters were used alongside the default augmented-coalescent likelihood.

### 2.5 Implementation

We have implemented the binary-state coalescent model as an open source R package `musseco` available at https://emvolz-phylodynamics.github.io/musseco. This includes a function, `fitbisseco`, that estimates multi-level selection coefficients by maximum likelihood and will estimate effective population size automatically if an estimate is not provided. The `fitbisseco` function requires a number of inputs in order to estimate the fitness parameters. A time-scaled phylogeny is required and can be estimated, for example, using the *treedater* R package [35]. This will also estimate the molecular clock rate of evolution, *µ*, which is another required input. Pathogen phylogenies are assumed to be reconstructed from population-based random samples of pathogen genomes and at most one sequence per host. The effective population size over time, *N*_*e*_(*t*), can also be input, although if it is not provided then it will be estimated with the *mlesky* R package [21] using the *skygrid* model [39] by default. This estimate of *N*_*e*_(*t*) is also provided in the returned BiSSECo fit using the `fitbisseco` function in the *musseco* R package [38]. Sample states, i.e. ancestral or variant, are also required to be defined for tips in the time-scaled phylogeny and included as an input to the `fitbisseco` function. An estimate of the generation time, *T*_*g*_, is also required. Initial estimates of *α, ω*, and *Y* (*t*) scaling parameter are also required as starting points for the estimation process, which is described in the Supporting Information.

Scripts used to simulate and analyse data are available at https://github.com/thednainus/musseco_simulations.

Scripts to analyse empirical data from HIV-1 are available at https://github.com/vinibfranc/BiSSECo_co-receptor_PANGEA.

## 3 Results

### 3.1 Simulations

Across most simulation studies, the augmented-coalescent outperformed the coalescent model that did not include a sampling likelihood based on mutation-selection balance. While the augmented-coalescent MLE generally had good precision (Fig 2A), the approximate confidence intervals did not always cover the true value, especially for birth-death simulations with high *R*_0_.

**Figure 2.**
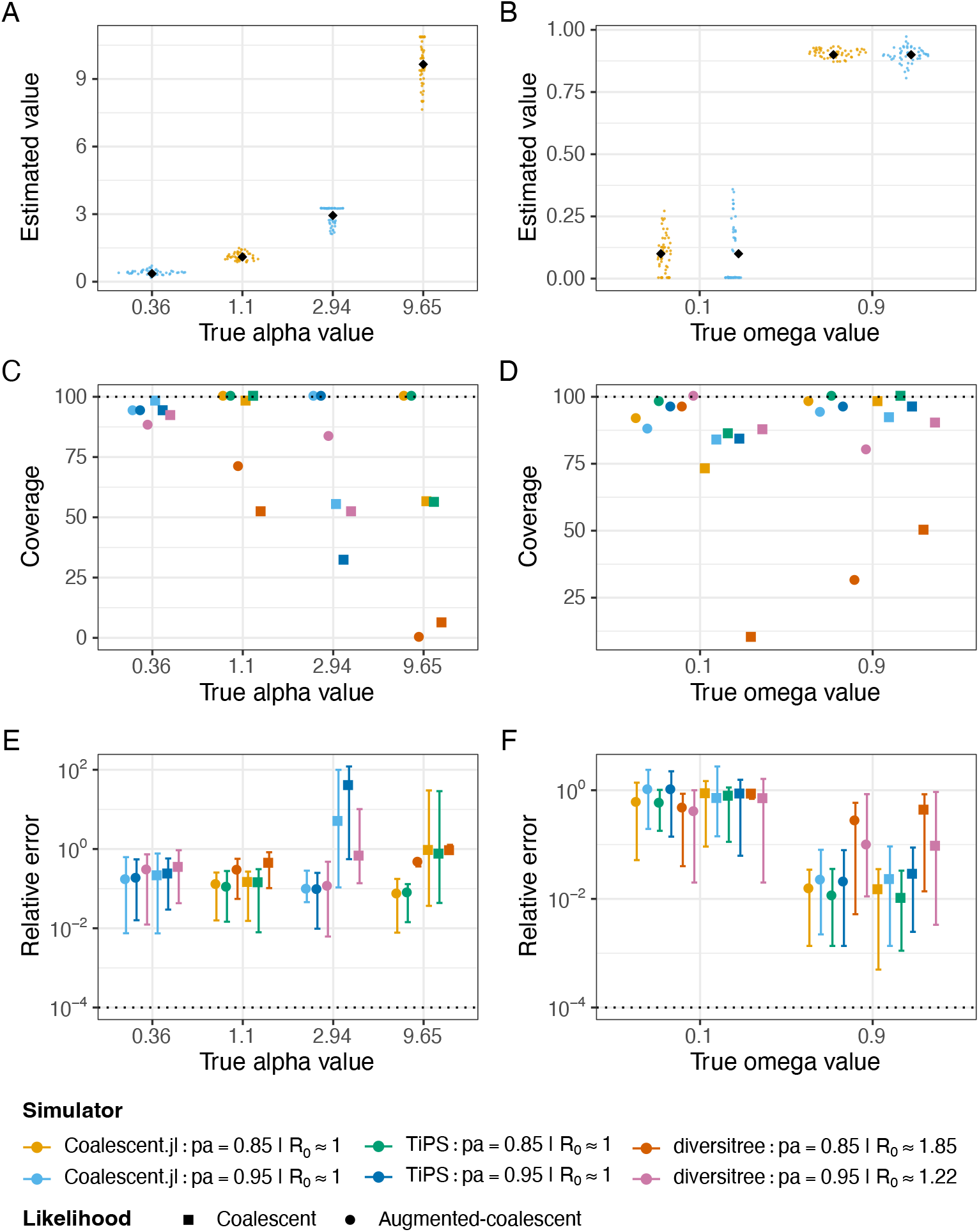
Simulation results for sample size of 1000 tips. (A) and (B) Sina plots showing the true and estimated values for *α* and *ω* for simulations carried out with *Coalescent*.*jl* and using the augmented-coalescent likelihood only. Black diamond shapes in the plot are the true values. (C) and (D) Coverage showing the percentage of replicates that contained the true *α* or *ω* values within the 95% credible interval for trees simulated with *Coalescent*.*jl, TiPS*, and *diversitree*. (E) and (F) Relative error for *α* and *ω* estimates. The relative error (y-axis) represents the computed median and the 0.025 and 0.975 quantiles for all replicates. Depending on the combination of parameter values, the proportion of individuals carrying the ancestral state (*p*_*a*_) could be 0.85 or 0.95. For the *diversitree* analyses, we grouped the values of *R*_0_ into two groups by averaging the observed *R*_0_ to 1.22 and 1.85 (see supporting information). For the *TiPS* /*Coalescent*.*jl* analyses, we sampled at equilibrium and *R*_0_ values are expected to be close to 1.0.

#### 3.1.1 Coverage

When estimating *α*, coverage was generally close to or equal to 100% for analyses carried out with *Coalescent*.*jl* or *TiPS*. However, when the true *α* value was 2.94 or 9.65, the augmented-coalescent likelihood performed better than the coalescent likelihood (Fig 2 and S5). For trees simulated with *diversitree*, it was clear that coverage was higher when *R*_0_ was low and the true value of *α* was 0.36 or 2.94. When *R*_0_ was high, coverage was close to zero when the true value of *α* was 9.65. In general, the augmented-coalescent likelihood also performed better than the coalescent likelihood (Fig 2 and S5).

When estimating *ω*, coverage was generally close to or equal to 100% when the true *ω* value was 0.9 for trees simulated using *Coalescent*.*jl* or *TiPS*. In most cases, the augmented-coalescent likelihood performed better than the coalescent likelihood (Fig 2 and S5). For analyses carried out with trees simulated with *diversitree*, we observed that coverage was close or equal to 100% when using the augmented-coalescent likelihood for the true *ω* value of 0.1 and independent of *R*_0_ or sample size (Fig 2 and S5). Coverage was lower when the true *ω* value was 0.9 (Fig 2 and S5). The augmented-coalescent likelihood was, in general, better when *ω* was small.

#### 3.1.2 Accuracy

For trees simulated with *Coalescent*.*jl* and *TiPS*, we observed that, in general, the estimates of *α* were very accurate showing lower values of relative error independent of sample size or true value of *α*. Note that when *α* was 2.94 or 9.65, the coalescent likelihood without augmentation produced very inaccurate results, showing relative error that was occasionally greater than 100-fold (Fig 2 and S5). For trees simulated with *diversitree*, accuracy was better for simulations in which the average *R*_0_ was 1.22 and independent of sample size. However, accuracy was worse for estimating values of *α* = 2.94 for lower *R*_0_ and using the coalescent likelihood, and this was fixed by using the augmented-coalescent likelihood (Fig 2 and S5).

For trees simulated with *Coalescent*.*jl* and *TiPS*, we observed that when *ω* was set to the low value of 0.1, accuracy remained low and was unaffected by sample size. In addition, the use of the augmented-coalescent likelihood did not substantially improve the estimates. However, when *ω* was set to a high value of 0.9, the estimates remained highly accurate and unaffected by sample size (Fig 2 and S5). For trees simulated with *diversitree*, accuracy was better when the average *R*_0_ was low. Furthermore, accuracy was slightly worse when *ω* was set to 0.1 (Fig 2 and S5).

Additional plots showing the estimated relative fitness parameters for individual replicates can be found in the supporting information (Fig S6 - S11). These figures span the range of tree simulators, numbers of tree tips, and both likelihood methods.

### 3.2 Simulation and analysis of sequence alignments

For analyses carried out with combination of parameters 1 (par1; Table S3), we were able to estimate both *α* and *ω* using sequence alignments of 2371 bp. The estimated time-scaled trees produced results similar to those obtained with the true trees (Fig 3). Furthermore, for par1 (Table S3), simulations were run for 45 years and the proportion of ancestral state reached 0.806 which was similar to the expected equilibrium value of 0.799 (Table S3).

**Figure 3.**
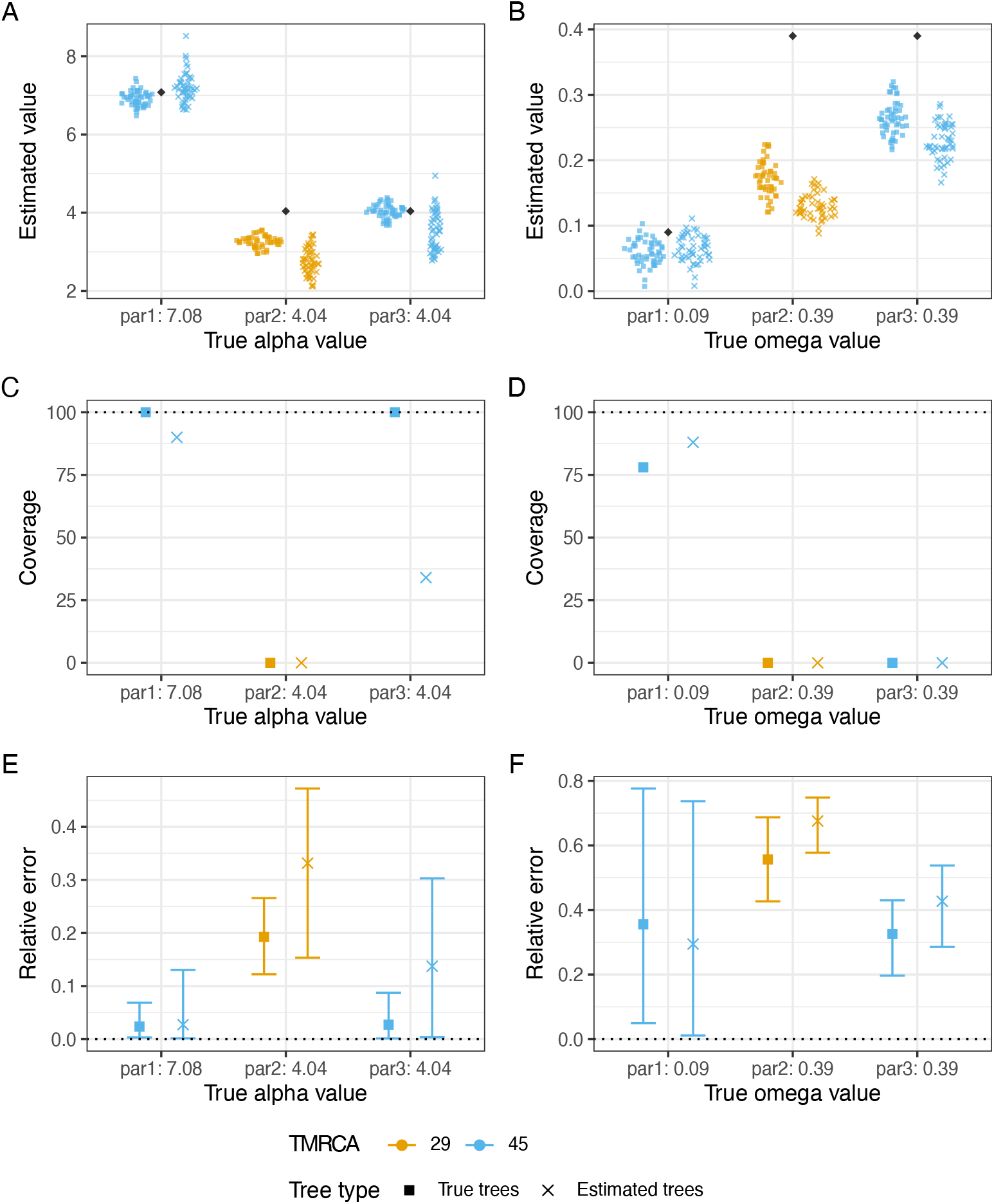
Performance in simulations with n=4,056. (A) and (B) Sina plots showing the true and estimated values for *α* and *ω* using the augmented-coalescent likelihood. Black diamond shapes in the plot are the true values. (C) and (D) Coverage showing the percentage of replicates that contained the true *α* or *ω* values within the 95% credible interval (E) and (F) Relative error for *α* and *ω* estimates. The relative error (y-axis) represents the computed median and the 0.025 and 0.975 quantiles for all replicates. All plots shows the estimated values for the true *Coalescent*.*jl* trees and estimated time-scaled trees. The par1, par2 and par3 refers to the combination of parameter values 1, 2 and 3 (Table S3). Sequence alignment length for par1 was 2371 bp and for par2 and par3 was 105 bp.

For analyses carried out with combinations of parameters 2 and 3 (par2 and par3; Table S3), we were unable to estimate *α* and *ω* using the true or estimated trees when the TMRCA (time to the most recent common ancestor) was 29 years. When the TMRCA was 45 years, we were able to estimate *α* using the true trees, but were unable to estimate *ω* with true or estimated trees (Fig 3). However, when the simulations were run for long enough and the state proportions reached equilibrium, we could estimate both relative fitness *α* and *ω* (Fig S4). Furthermore, for par2 and par3 (Table S3), the proportion of ancestral state was 0.865 and 0.822, respectively, suggesting that it did not reach equilibrium (Table S3). We were also unable to recover the true TMRCA and substitution rates with *treedater* using trees reconstructed with alignments of 105 bp (Fig S3).

### 3.3 HIV-1 co-receptor tropism

The co-receptor tropism prediction of the subtype C PopART Phylogenetics sequences from Zambia revealed that 20.0% are capable of using the X4 co-receptor, associated with faster disease progression. This analysis showed that: (i) WebPSSM scores consistently split R5- and X4-capable viruses, (ii) there was a slight increase in the proportion of X4/R5×4 and decrease of R5 over the study period, (iii) the pre-treatment CD4 counts were similar for individuals infected by R5 and X4-capable viruses, and (iv) the proportion of X4/R5×4 tends to be higher for individuals in advanced stages of infection (low CD4) when compared with earlier stages (Fig S12).

As expected given high within-host selective pressures for co-receptor switching, the fold-change in within-host replicative fitness (*α*) was much higher than one (6.87, 95% CI: 6.08–7.76) for the *env* analysis, while the fold-change in between-host replicative fitness (*ω*) was much lower than one (0.09, 95% CI: 0.04–0.18). These values are consistent with the hypothesis that the within-host relative fitness of variant (X4/R5×4) is substantially higher than the ancestral type (R5), while the between-host fitness of X4/R5×4 is much lower than R5. In other words, the magnitude of the transmission penalty of X4-capable viruses is approximately −90%.

## 4 Discussion

We have developed a structured coalescent model which describes pathogen evolution of two co-circulating variants with multi-level (within- and between-host) selection. We further develop a maximum-likelihood method for inferring multi-level selection coefficients based on this model and a convenient software implementation. This approach builds on previous work on related structured birth-death models (e.g. the BiSSE model) and adds new capabilities: This model accounts for nonlinear dynamics in effective population size, variable sampling through time, and provides a novel integration of a sampling likelihood with the genealogical coalescent likelihood. In simulations, we find that the derived mutation-selection balance used in combination with the coalescent likelihood provides superior estimates of multi-level selection parameters. We further demonstrate that within-host selection coefficients can be estimated from population-based samples across a large range of values, from minor fitness costs to highly deleterious phenotypes. This provides an original avenue for estimating fitness cost of common deleterious variants from epidemiological data which has not traditionally been used for this purpose. While our focus has been on evolution of infectious disease pathogens, the computational framework is quite general, and could also be applied to study macro-evolution.

In our simulations, we also demonstrated that when the genetic sequence alignment was sufficiently large and informative (similar to the empirical *env* gene), and the proportion of ancestral state and variant had reached equilibrium, we were able to estimate the relative fitness *ω* and *α* (Fig 3). Our empirical analysis of the evolution of HIV co-receptor tropism supports the commonly hypothesised reduction in transmission associated with the X4-tropic virus and contra-indicates the “random transmission” hypothesis [10, 11] according to which both X4 and R5 tropic virus are transmitted equally in proportion to their viral titre. Our study appears to be unique in estimating a specific value for the fitness cost. This analysis, performed using data from a real and high HIV prevalence setting, confirmed that, when compared to R5, the within-host replicative fitness (*α*) of X4/R5×4 variants was higher (seven-fold increase) and the between-host fitness (*ω*) was much lower (≈ −90%). This estimate suggests that there are selective forces acting against X4 transmission and provides a magnitude for this transmission penalty [8, 9]. Therefore, the lower frequency of X4 viruses in genital secretions is not the only factor acting against their transmission, although it is still very possible that X4/R5×4 variants will be transmitted if they dominate the donor’s virus population, consistent with X4-tropic virus being transmitted at 9% of the rate of R5-tropic virus. The *α* estimates demonstrate pronounced within-host selection acting on X4/R5×4.

There are several limitations in our study. First, we used a single value for the infection generation time. However, it is possible that for the co-receptor analysis, infections with different co-receptors may lead to different rates of disease progression. Second, current state-of-the-art bioinformatic methods for the classification of co-receptor usage do not distinguish between the dual-receptor R5×4 from the single receptor X4. Consequently, we were unable to test whether dual receptor usage would provide any advantage in replicative fitness compared to X4 usage only. There is potential for this method to be expanded to incorporate more than two competing variants, such as adaptation driven by differing human leukocyte antigen (HLA) types in a host population. For example, the methods used by Palmer et al. [40] could be used to identify nucleotide sites that are linked to evolutionary pressure from HLA type and treatments that lead to drug resistance mutations (DRM), thereby allowing the definition of multiple phenotypes for analysis using the model presented here. While in its current form, we expect there to be an issue with statistical-identifiability as the number of parameters in the model is increased, this is an area for future development. Other developments may be possible related to the incorporation of more than one sequence per host, which if available could greatly enhance within-host selection estimates, as well as the incorporation of additional metadata (e.g. structural data or replicative capacity assays) when estimating selection coefficients.

## 6 Acknowledgements

The authors thank the Phylogenetics and Networks for Generalised HIV Epidemics in Africa (PANGEA) consortium for access to the PopART Zambian HIV-1 dataset. PANGEA is funded by the Bill & Melinda Gates Foundation (consecutive grants OPP1084362 and OPP1175094).

## PANGEA consortium members

Christophe Fraser^1^, Kate Grabowski^2^, Sikhulile Moyo^3^, Deenan Pillay^4^, Andrew Rambaut^5^, Oliver Ratmann^6^, Deogratius Ssemwanga^7^, Lucie Abeler-Dörner^1^, Helen Ayles^8^, David Bonsall^1^, Rory Bowden^1^, Max Essex^9^, Sarah Fidler^10^, Tanya Golubchik^1^, Ravindra Gupta^11^, Richard Hayes^12^, Joshua Herbeck^13^, Joseph Kagaayi^14^, Pontiano Kaleebu^7^, Jairam Lingappa^13^, Vladimir Novitsky^15^, Thomas Quinn^2^, Janet Seeley^7,16^, Frank Tanser^17^, Maria Wawer^2^, Myron Cohen^18^, Vincent Calvez^19^, Thumbi Ndung’u^17^, Tulio D’Oliveira^20^, Ann Dennis^18^, Dan Frampton^4^, Anne Hoppe^4^, Paul Kellam^11^, Cissy Kityo^21^, Andrew Leigh-Brown^5^, Nick Paton^22^.

^1^University of Oxford, Oxford, UK, ^2^Johns Hopkins University, Baltimore, USA, ^3^Botswana Harvard AIDS Institute Partnership, Botswana, ^4^University College London, London, UK, ^5^University of Edinburgh, Edinburgh, UK, ^6^Imperial College London, London, UK, ^7^Medical Research Council/Uganda Virus Research Institute, Entebbe, Uganda, ^8^PopART/Zambart, Lusaka, Zambia, ^9^Harvard Botswana, Gaborone, Botswana, ^10^PopART/Imperial College London, UK, ^11^University of Cambridge, Cambridge, UK, ^12^PopART/London School of Hygiene and Tropical Medicine, London, UK, ^13^University of Washington, Seattle, USA, ^14^Rakai Health Sciences Program, Kalisizo, Uganda, ^15^Harvard University, Boston, USA ^16^London School of Hygiene and Tropical Medicine, London, UK, ^17^Africa Health Research Institute, South Africa, ^18^University of North Carolina, USA, ^19^Institut Pasteur, Paris, FR, ^20^University of KwaZulu-Natal, South Africa, ^21^EARNEST/The Joint Clinical Research Centre, Kampala, Uganda, ^22^EARNEST/University of Singapore, Singapore.

## Supporting information captions

### S1 Text. Supporting Information

#### Data availability statement

The method used here is available as an open source R package *musseco* at https://github.com/emvolz-phylodynamics/musseco and there is a tutorial at https://emvolz-phylodynamics.github.io/musseco. Scripts used to simulate and analyse data are available at https://github.com/thednainus/musseco_simulations. Scripts to analyse empirical data from HIV-1 are available at https://github.com/vinibfranc/BiSSECo_co-receptor_PANGEA. Access to the PANGEA PopART HIV-1 sequence dataset is through application. Details are available at https://www.pangea-hiv.org/join-us.

