## Supporting Information for "Untangling the effects of within-host and between-host natural selection in viruses with applications to evolution of HIV-1 co-receptor usage"

Short title: Quantifying within-host and between-host selection in virus evolution

Kieran O. Drake[1¶], Vinicius B. Franceschi[1¶], Fabrícia F. Nascimento[1¶], Erik M. Volz[1¶*], on behalf of the PANGEA Consortium[†]

[1]MRC Centre for Global Infectious Disease Analysis, School of Public Health, Imperial College London, London, UK

[†]A full list of the PANGEA consortium members appear at the end of the manuscript

[¶]These authors contributed equally to this work.

28 April, 2025

### Supporting Information

#### 1 Methods

##### 1.1 The binary-state speciation and extinction coalescent

In a general compartmental model, such as the common susceptible-infected-removed (SIR) model [1], neither the effective population size nor migration rate between demes need be constant, and in more general frameworks, coalescence is also allowed between lineages occupying different demes [2].

Here we use a coalescent model to describe a deterministic non-linear dynamical system and thereby the epidemiological and demographic history. Using this we can derive and compute the log-likelihood of a genealogy with heterochronous sampling and labeled taxa. By computing the log-likelihood for a range of parameter values we can obtain a maximum log-likelihood estimate for those parameters and thereby infer the 'true' values. In this study we apply this method to untangle the effects of within-host and between-host selection in virus evolution. The specific parameters that we have investigated are the relative rates for transmission and mutation for two variants.

The *musseco* R package [3] achieves this by taking input parameters for the molecular clock rate of evolution, $\mu$, the generation time, $T_g$, (i.e. the mean time elapsed between generations), a time-scaled phylogeny, and an estimate of the time-varying effective population size, $N_e(t)$, although if the latter is not provided then it will be automatically estimated by the package. An epidemiological model is defined and a demographic history is simulated for the populations in each deme within the model. The log-likelihood that this modeled demographic history results in the input time-scaled phylogeny is then computed for a given set of parameters. This is repeated using an optimisation algorithm to find the maximum log-likelihood. The fitness parameter values used in the model identified as having the maximum log-likelihood in relation to the time-scaled phylogeny are our inferred parameter values. The optimisation algorithm also enables us to compute confidence intervals around the maximum log-likelihood estimates for the inferred fitness parameters. This workflow is summarised in Figure S1 and described in further detail in the following sections.

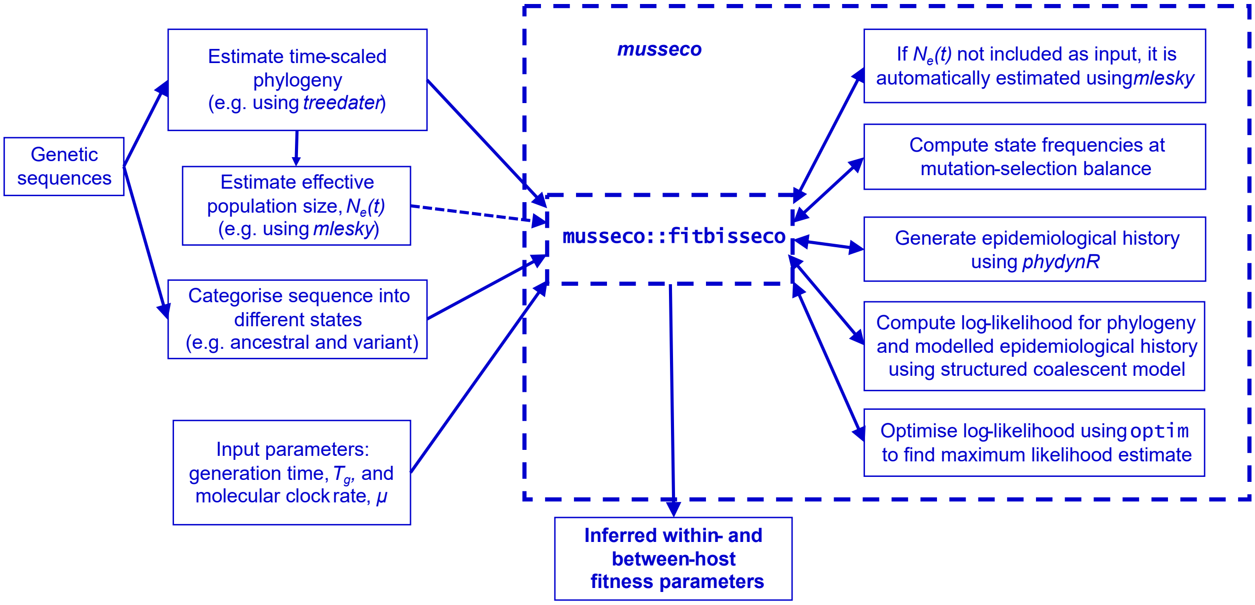


Figure S1: Workflow for estimating relative fitness of variants using the *musseco* R package. Using a coalescent likelihood approach, the `fitbisseco` function included in the package can infer within-host ($\alpha$) and between-host ($\omega$) relative fitness parameters from a time-scaled phylogeny labeled with variant states.

##### 1.1.1 Coalescent likelihood conditional on epidemiological history

The structured coalescent incorporates population structure by extending basic coalescent models, originally developed by Kingman [4, 5], to allow lineages to migrate between different subpopulations or demes [6]. While the structured coalescent is often used to incorporate geographic population structure, the theory holds for many different forms of population structure [7, 8]. In this work, our demes represent phylotypes, ancestral and variant, rather than geographic locations. It is worth noting therefore that 'migration' and 'mutation' are interchangeable in this context as they both indicate a change of deme.

A variety of approximations have been developed for computing coalescent likelihoods [9]. We use the fast likelihood 'PL2' approximation developed by Volz & Siveroni [2], which is included in the *PhyDynR* package [10]. In common with other structured coalescent models, PL2 defines a discrete Markovian stochastic tree-building process in retrospective/continuous time [7] and defines the potentially time and state dependent rates that a lineage changes demes (e.g. from deme $k$ to $l$ at rate $M_{kl}(t)$) and the rate of coalescence in which two or more lineages merge to form a single lineage (e.g. a lineage in deme $k$ 'gives birth' to a lineage in deme $l$ at rate $\phi_{kl}(t)$). The

PL2 model has improved accuracy relative to previous iterations [11, 12], although this comes at a higher computational cost. Similar to the earlier models, it works by solving a system of differential equations for each lineage, but it better accounts for correlations between co-existing lineages and effects stemming from disparate coalescent rates between demes.

The lineage-deme configuration through time, $S(t)$, is only observed at times of sampling and estimation of migration rates and/or effective population sizes, $Ne(t)$, is typically made with a Markov chain Monte Carlo (MCMC) simulation to marginalize over latent $S(t)$ [13]. However, this entails a large computational cost of integrating over a very large configuration space [14]. To reduce the computational cost, Volz & Siveroni [2], marginalize over lineage states, deriving an approximate likelihood in terms of the probability that a lineage is in a particular deme. After making the simplifying approximation that the conditional probability of state $j$ is equal to the marginal, $p(s_j = l|s_i = k) \approx p(s_j = l)$, they show that rates of deme-change for lineage $i$ and the rate of coalescence of $i$ with all other lineages can be computed. Where the configuration is unknown, the total rate of coalescence, $\lambda_{ij}$, between lineages $i$ and $j$ can then be approximated by equation 1:

$$\lambda_{ij}(t) = p_i'\Phi p_j + p_j'\Phi p_i, \tag{1}$$

where $p_i$, $p_i'$, $p_j$, $p_j'$ represent the probabilities of lineages being in different demes and $\Phi$ is the rate at a which a lineage in deme $l$ 'gives birth' to a lineage in deme $k$.

An approximation for the cumulative hazard of coalescence between consecutive time points for lineage $i$, $\Omega_i$, can be shown to be:

$$\Omega_i \approx \int_{t=t_i}^{t_i+1} \sum_{i\in\mathcal{A}(t)} \sum_{\substack{j\in\mathcal{A}(t)\\ j\neq i}} \lambda_{ij}(t)/2\,dt, \tag{2}$$

where $\mathcal{A}(t)$ represents the set of extant lineages. By using this approximation Volz & Siveroni [2] show that we can avoid computing the full intractable probability density of the genealogy configuration at all time points.

Equation 3 is an approximation of the time dependent migration matrix for lineage $i$ and is a function of migration events, $G_{lk}(t)$, and birth events, $F_{lk}(t)$, which change the deme of a lineage.

As before, this relies on the approximation that $p_{jl}(t) \approx p(s_j(t) = l | s_i(t) = k)$ because the joint density for $s_i$ and $s_j$ is not tractable.

$$M^{(i)}_{kl}(t) \approx \begin{cases} \left(G_{lk}(t)+F_{lk}(t)(1-\frac{1}{Y_t(t)}\sum_{j\in\mathcal{A}(t)} p_{jl}(t))\right)/Y_k(t) & \text{if } k\neq l \\ 0 & \text{if } k=l \end{cases} \tag{3}$$

To account for the effect of coalescence on evolution of the state vector $p_i(t)$, Volz & Siveroni [2] instead define the augmented state vector, $\tilde{p}_i(t)$, which has $m + 1$ elements and where $\tilde{p}_{i,m+1}(t)$ represents the probability that lineage $i$ has coalesced at some time $t' < t$. The forward equations for the evolution of $\tilde{p}_i$ can then be found by tabulating the effects of migration, birth, and coalescent events, as shown in equation 4.

$$\begin{aligned} \tilde{p}_{ik}(t + \Delta t)|k < m{+}1 = \tilde{p}_{ik}(t) & \\ + (\Delta t) \sum_{l\neq k} M^{(i)}_{lk} \tilde{p}_{il}(t) - M^{(i)}_{kl} \tilde{p}_{ik}(t) & \qquad \text{“Migration”} \\ - (\Delta t) \sum_{i}^{m} \tilde{p}_{ik}(t)(\phi_{kl}(t) + \phi_{lk}(t))(A_l(t) - p_{il}(t)) & \qquad \text{“Coalescent”} \\ + \mathcal{O}(\Delta t) & \end{aligned} \tag{4}$$

By computing the limit $\Delta t \to 0$ and making appropriate substitutions of $M(t)$ and $\phi(t)$ for $G(t)$ and $F(t)$, as shown in equations 3 and 4 in the main text, a system of differential equations can be defined. Note that $\mathcal{O}(\Delta t)$ represents the terms that tend to zero faster than $\Delta t$. This system of differential equations can be solved for each lineage and minimised to find the maximum log-likelihood solution and the corresponding rates of migration and coalescence.

In the study presented here, these represent the mutation rates, $\mu_{av}$ and $\mu$, and transmission rates, $\beta_a$ and $\beta_v$, as shown in equations 2 and 1 in the main text.

The input parameters to the log-likelihood function are optimised to result in the maximum log-likelihood estimate using the `optim` function in the *stats* R package. The parameters resulting in the maximum log-likelihood estimate enable us to infer the values of $\alpha$ (within-host relative fitness) and $\omega$ (between-host relative fitness). The results output by the package also include 95% confidence interval values as computed using the Fisher information method. More precisely, we use the standard errors of the negative Hessian matrix, as computed within the `optim` function in the *stats* R package, which approximates the Fisher information matrix at the optimum.

There are limitations to this approach to confidence interval estimation, which mean that when the likelihood surface is flat or close to flat in the region of the maximum value, i.e. when there is low Fisher information, it may not return reasonable confidence interval bounds, if indeed an upper boundary exists. This effect is discussed in section 2.1.2. Computing confidence intervals using a profile likelihood method may improve this for some parameter sets but at higher computational cost and some loss of speed so it has not been implemented here.

###### 1.1.2 Inference by maximum likelihood estimate

We use a maximum log-likelihood approach to infer the relative fitness parameters of interest: within-host relative fitness, $\alpha$, and between-host relative fitness, $\omega$, as defined previously. A scaling factor, $\iota$, for $Y(t)$, the total number of infected hosts, is also inferred using this method.

There are two maximum log-likelihood (MLL) methods available within the *musseco* package. Both methods are based on the conditional density of a genealogy given epidemic and demographic parameters as implemented in the PL2 structured coalescent likelihood method as summarised in section 1.1.1. We name the first of these two MLL methods the 'coalescent' likelihood. This relies solely on the independently estimated time-scaled phylogeny input and the demographic history defined by the model to infer the fitness parameters. The second, we name the 'augmented-coalescent' likelihood. This combines, in a simple sum, the coalescent likelihood with a binomial likelihood of sampling variants or ancestral types under the assumption of random sampling and mutation-selection balance. The augmented-coalescent likelihood is the default option in *musseco*. The relative performance of the 'coalescent' and 'augmented-coalescent' likelihood methods is discussed in the Results section in the main text.

##### 1.2 Implementation

The `fitbisseco` function will run with default values for $\alpha$, $\omega$, and the $Y(t)$ scaling parameter equal to 15, 0.95, and 1 respectively unless they are changed within the '*theta0*' argument. Likewise, if a time series for the effective population size, $N_e(t)$, is not provided then it will be estimated using the default parameters, although these can be changed using the '*mlesky_parms*' argument.

#### 1.3 Simulations

The parameter values used to simulate trees with *Coalescent.jl* and *TiPS* are listed in Table S1. The parameter values used to simulate trees with *diversitree* are described in Table S2. For *diversitree* simulations, we used a similar nomenclature as those described in [15], but using terms related to epidemiology. For example, instead of speciation and extinction rates as described in [15], we used birth and mortality (removal) rates, respectively.

Table S1: Parameter values used to simulate genealogies using *Coalescent.jl* and *TiPS*

| **Definition and Symbol** | **Fixed value** | | | |
|---|---|---|---|---|
| Initial numbers of infectious $Y_a$ individuals | 1 | 1 | 1 | 1 |
| Initial numbers of infectious $Y_v$ individuals | 0 | 0 | 0 | 0 |
| Growth rate ($\beta$) | 0.216 | 0.216 | 0.216 | 0.216 |
| Carrying capacity ($K$) | 10,000 | 10,000 | 10,000 | 10,000 |
| Mortality rate ($\gamma$)[a] | 1/10.2 | 1/10.2 | 1/10.2 | 1/10.2 |
| Transition from $Y_v$ to $Y_a$ ($\mu_{va}$)[b] | 0.0016 | 0.0016 | 0.0016 | 0.0016 |
| Transition from $Y_a$ to $Y_v$ ($\mu_{av}$) | 0.0018 | 0.015 | 0.0006 | 0.005 |
| Selection coefficient ($s$) | -0.1 | -0.9 | -0.1 | -0.9 |
| Between-host relative fitness ($\omega$)[c] | 0.9 | 0.1 | 0.9 | 0.1 |
| Within-host relative fitness ($\alpha$)[d] | 1.10 | 9.65 | 0.36 | 2.93 |
| Proportion of individuals in ancestral state[e] | 0.85 | 0.85 | 0.95 | 0.95 |

[a] Based on [16] who estimated the median survival time in Africa as 10.2 years when considering individuals 30 years old at seroconversion.
[b] Based on estimates obtained for HIV-1 subtype C [17].
[c] $\omega = 1 + s$
[d] $\alpha = \mu_{av}/\mu$
[e] at equilibrium

Table S2: Parameter values used to simulate genealogies using *diversitree*

| **Definition and Symbol**[a] | **Fixed value** | | | |
|---|---|---|---|---|
| Birth rate of state 0 ($\lambda_0$) | 0.1800 | 0.0200 | 0.1125 | 0.0125 |
| Birth rate of state 1 ($\lambda_1$) | 0.2000 | 0.2000 | 0.1250 | 0.1250 |
| Mortality rate of state 0 ($\mu_0$) | 0.1000 | 0.1000 | 0.1000 | 0.1000 |
| Mortality rate of state 1 ($\mu_1$) | 0.1000 | 0.1000 | 0.1000 | 0.1000 |
| Transition rate from 0 to 1 ($q_{01}$) | 0.0016 | 0.0016 | 0.0016 | 0.0016 |
| Transition rate from 1 to 0 ($q_{10}$) | 0.0018 | 0.015 | 0.0006 | 0.005 |
| Within-host relative fitness ($\alpha$)[b] | 1.1 | 9.65 | 0.36 | 2.93 |
| Between-host relative fitness ($\omega$)[c] | 0.9 | 0.1 | 0.9 | 0.1 |
| Basic reproduction number ($R_0$) | 1.97 | 1.74 | 1.24 | 1.19 |
| Proportion of individuals at state 1[d] | 0.85 | 0.85 | 0.95 | 0.95 |

[a] For simulating trees with the BiSSE model implemented in the *diversitree* R package [15], we defined states 0 and 1, that here represents the variant and ancestral states, respectively.
[b] $\alpha = q_{10}/q_{01}$
[c] $\omega = \lambda_0/\lambda_1$
[d] at equilibrium

#### 1.4 Simulation and analyses of genetic sequence alignments

We used the program *Coalescent.jl* to simulate genealogies composed of 4056 tips. The parameter values used to simulate these trees are described in Table S3.

Table S3: Parameter (par) values used to simulate genealogies using *Coalescent.jl* to mimic observed HIV data.

| **Definition and Symbol** | **par1**[a] | **par2**[b] | **par3**[c] |
|---|---|---|---|
| Initial numbers of infectious $Y_a$ individuals | 1 | 1 | 1 |
| Initial numbers of infectious $Y_v$ individuals | 0 | 0 | 0 |
| Growth rate ($\beta$) | 0.863 | 0.863 | 0.863 |
| Carrying capacity ($K$) | 10,000 | 10,000 | 10,000 |
| Mortality rate ($\gamma$)[d] | 1/10.2 | 1/10.2 | 1/10.2 |
| Transition from $Y_v$ to $Y_a$ ($\mu_{va}$) | 0.0032 | 0.0036[c] | 0.0036[c] |
| Transition from $Y_a$ to $Y_v$ ($\mu_{av}$) | 0.0226 | 0.0145 | 0.0145 |
| Selection coefficient ($s$) | -0.91 | -0.61 | -0.61[c] |
| Between-host relative fitness ($\omega$)[d] | 0.09 | 0.39[c] | 0.39 |
| Within-host relative fitness ($\alpha$)[e] | 7.07 | 4.04 | 4.04 |
| Proportion of individuals at ancestral state[f] | 0.79 | 0.79 | 0.79 |
| TMRCA[g] | 45 | 29 | 45 |

[a] Observed value when analysing the sequence alignment for the HIV-1 *env* gene.
[b] Observed value when analysing the sequence alignment for the HIV-1 V3 loop.
[c] Observed value when analysing the sequence alignment for the HIV-1 V3 loop, but using the same TMRCA as observed for the *env* gene.
[d] Based on [16] who estimated the median survival time in Africa as 10.2 years when considering individuals 30 years old at seroconversion.
[d] $\omega = 1 + s$
[e] $\alpha = \mu_{av}/\mu$
[f] at equilibrium
[g] Time to the most recent common ancestor in years.

Alignments of 105 bp (after removal of gaps observed in most sequences) for the V3 loop and 2371 bp for the *env* gene as observed for the empirical HIV-1 data were simulated with *AliSim* [18]. We simulated these alignments using the GTR+F+I substitution model. This substitution model was used with the empirical V3 loop and *env* to estimate maximum likelihood (ML) trees with `IQ-TREE` [19]. `IQ-TREE` esimated the following values for (1) V3 loop: GTR3.51951, 7.79129, 2.15558, 3.12071, 21.10303, 1.00000 +F0.444, 0.156, 0.220, 0.180 +I0.063 and (2) *env* gene: GTR2.21430, 5.04449, 0.73827, 0.72478, 5.39684, 1.0 +F0.342, 0.169, 0.246, 0.244 +I0.117. These were the values used to simulate the genetic sequence alignments.

For each of the empirical V3 loop and *env* alignments, a time-scaled tree was estimated with the *treedater* R package [20]. We obtained a mean substitution rate of 0.0036 and 0.0032 for V3 loop and *env*, respectively. Those values were also used to simulate sequence alignments with *AliSim*.

### 2 Results

#### 2.1 Simulations

##### 2.1.1 Individual replicates

We plotted the confidence interval for all replicates analysed in this study. These plots are available at `https://github.com/thednainus/musseco_simulations/tree/main/plots_paper/supporting_information/individual_replicates`.

##### 2.1.2 Upper bound of the confidence interval

We calculated the percentage of replicates in which we were unable to compute the upper bound of the confidence interval (CI) using the Fisher information matrix at the optimum (see Methods in the main text and section 1.1.1 above).

For trees simulated with *Coalescent.jl* and *TiPS*, we observed that for approximately 25–35% of the replicates in which the true value of $\alpha$ was 2.94 or 9.65, we could not compute the upper bound of the CI using the coalescent likelihood. This reached >60% for trees simulated with *Coalescent.jl* with sample size of 500 individuals and also using the coalescent likelihood (Fig S2).

For trees simulated with *diversitree*, when $R_0$ approximately 1.22, for approximately 25% of replicates we could not compute the upper bound when using the coalescent likelihood (Fig S2).

For trees simulated with *Coalescent.jl* and *TiPS*, we observed that we could not compute the upper bound of the CI for approximately 60% of the replicates when the true value of $\omega$ was low using both coalescent and augmented-coalescent likelihoods. This was particularly worse for sample size of 500 individuals (Fig S2).

For trees simulated with *diversitree*, we observed, in general, a lower percentage (approximately 8%) of replicates in which we could not compute the upper bound of the CI for the lower $\omega = 0.1$ and when using the coalescent or augmented-coalescent likelihoods and using 1000 or 500 tips (Fig S2). However, for the case in which the $R_0$ was approximately 1.22, the percentage of replicates that we could not compute the upper bound was higher at approximately 30%.

Finally, for all trees simulated with 4,056 tips using the coalescent-augmented likelihood, we were always able to compute the upper bound of the confidence interval.

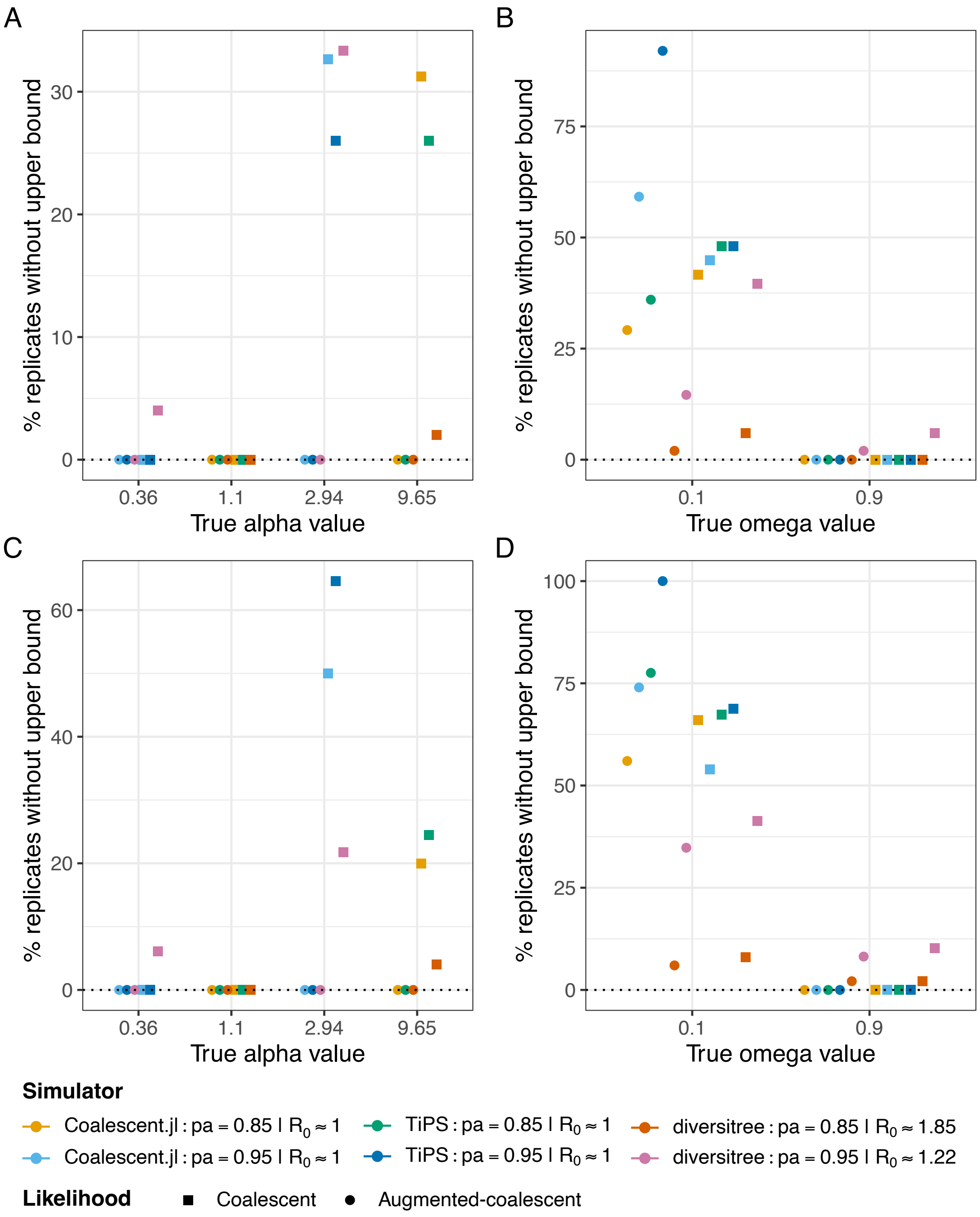


Figure S2: **Upper bounds**: Percentage of replicates in which the upper bound for the 95% credible interval could not be computed for trees simulated with *Coalescent.jl*, *TiPS* and *diversitree* and for sample sizes of (A and B) 1000 and (C and D) 500 individuals. Depending on combination of parameter values, the proportion of individuals carrying the ancestral state ($p_a$) could be 0.85 or 0.95. For the *diversitree* analyses, we grouped the values of $R_0$ into two groups by averaging the observed $R_0$ to 1.22 and 1.85 (see Table S2). For the *TiPS*/*Coalescent.jl* analyses, we sampled at equilibrium and $R_0$ are expected to be close to 1.0. x-axis shows the true value of (A and C) $\alpha$ and (B and D) $\omega$.

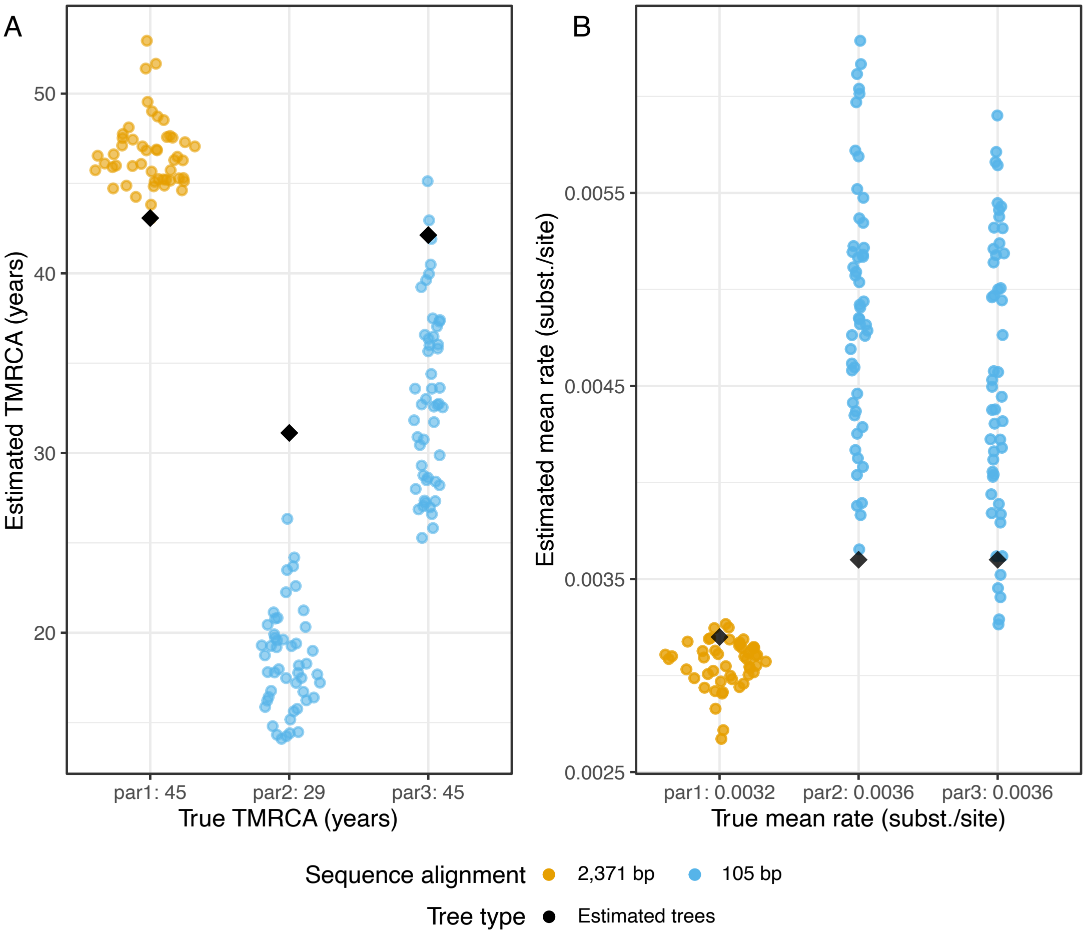


Figure S3: **Simulation results for sample size of 4,056 tips**: Sina plots showing the true and estimated values for the (A) TMRCA and (B) mean substitution rate for the time-scaled trees using alignments of 2371 bp and 105 bp. For parameter values (par1, par2 and par3) see Table S3.

#### 2.2 Simulation and analyses of genetic sequence alignments

When using alignments of 2371 bp, *treedater* was able to correctly estimate the TMRCA (time to the most recent common ancestor) and the mean substitution rate used in our simulations (Fig S3). However, when using alignments of 105 bp, *treedater* was unable to recover the TMRCA and the mean substitution rate (Fig S3).

We also carried out an additional simulation for the parameter values observed for the empirical V3 loop, but running long enough (TMRCA = 1000 years) to allow the proportion of ancestral and variant states to reach equilibrium. For these analyses, we were able to estimate both $\alpha$ and $\omega$ (Fig S4).

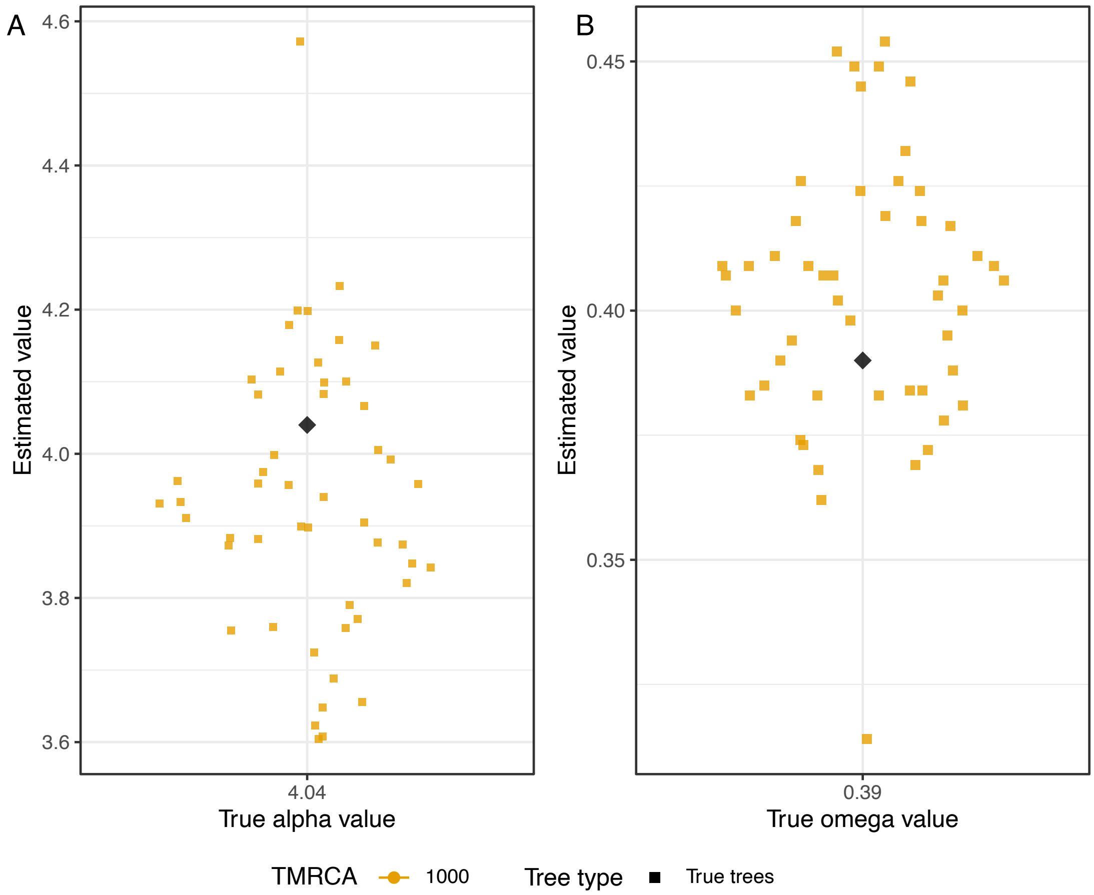


Figure S4: **Precision**: Sina plots showing the true and estimated values for the (A) $\alpha$ and (B) $\omega$ for the true *Coalescent.jl* trees using the combination of parameter values obtained for the empirical V3 loop and TMRCA = 1000 years.

#### 3 Additional supporting figures and tables

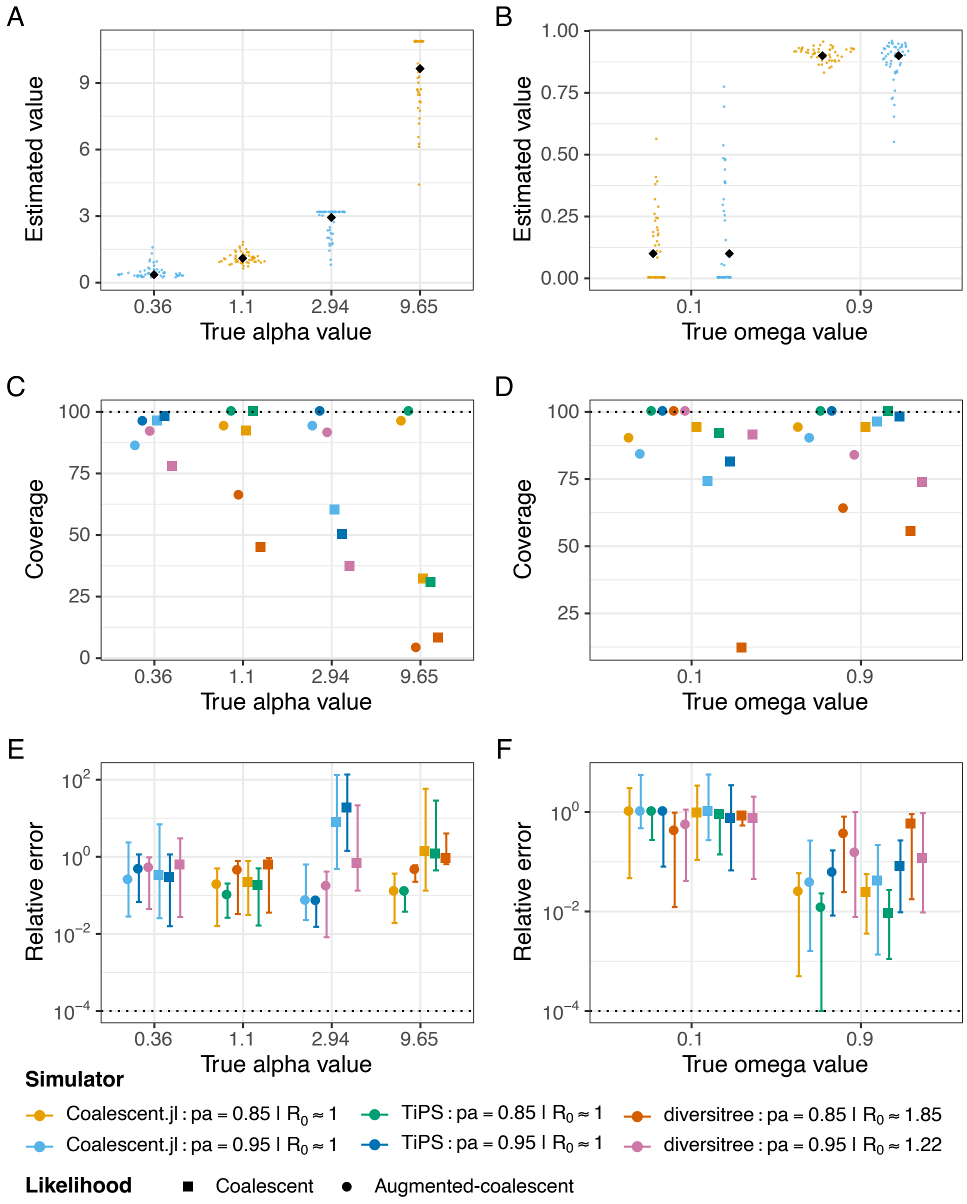


Figure S5: **Simulation results for sample size of 500 tips**: (A) and (B) Sina plots showing the true and estimated values for $\alpha$ and $\omega$ for simulations carried out with *Coalescent.jl* and using the augmented-coalescent likelihood only. Black diamond shapes in the plot are the true values. (C) and (D) Coverage showing the percentage of replicates that contained the true $\alpha$ or $\omega$ values within the 95% credible interval for trees simulated with *Coalescent.jl*, *TiPS*, and *diversitree*. (E) and (F) Relative error for $\alpha$ and $\omega$ estimates. The relative error (y-axis) represents the computed median and the 0.025 and 0.975 quantiles for all replicates. Depending on the combination of parameter values, the proportion of individuals carrying the ancestral state ($p_a$) could be 0.85 or 0.95. For the *diversitree* analyses, we grouped the values of $R_0$ into two groups by averaging the observed $R_0$ to 1.22 and 1.85 (see Table S2). For the *TiPS*/*Coalescent.jl* analyses, we sampled at equilibrium and $R_0$ is expected to be close to 1.0.

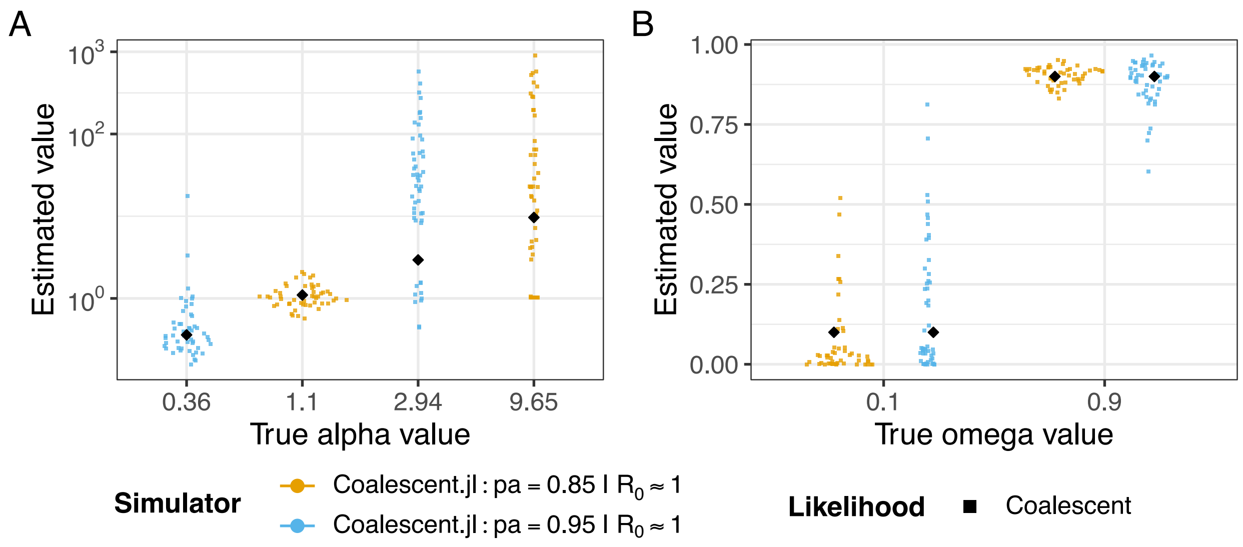


Figure S6: **Sina plot for *Coalescent.jl* simulations with 500 tips**: True versus estimated (A) $\alpha$ and (B) $\omega$ values using the coalescent likelihood only. Black diamond shapes in the plot are the true values. Depending on combination of parameter values, the proportion of individuals carrying the ancestral state ($p_a$) could be 0.85 or 0.95. For these simulations, we sampled at equilibrium and $R_0$ is expected to be close to 1.0.

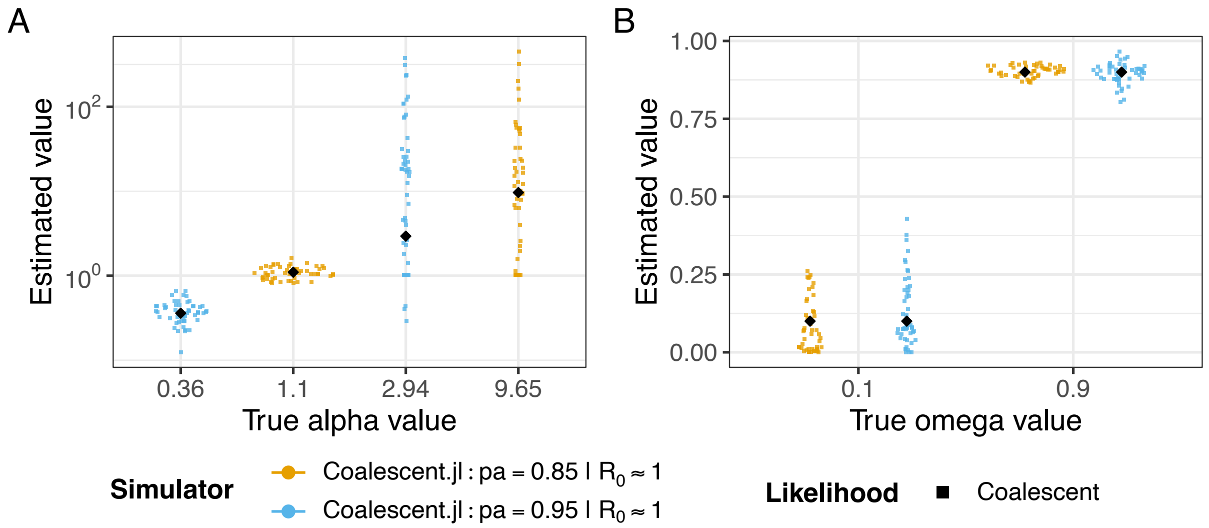


Figure S7: **Sina plot for *Coalescent.jl* simulations with 1000 tips**: True versus estimated (A) $\alpha$ and (B) $\omega$ values using the coalescent likelihood only. Black diamond shapes in the plot are the true values. Depending on combination of parameter values, the proportion of individuals carrying the ancestral state ($p_a$) could be 0.85 or 0.95. For these simulations, we sampled at equilibrium and $R_0$ is expected to be close to 1.0.

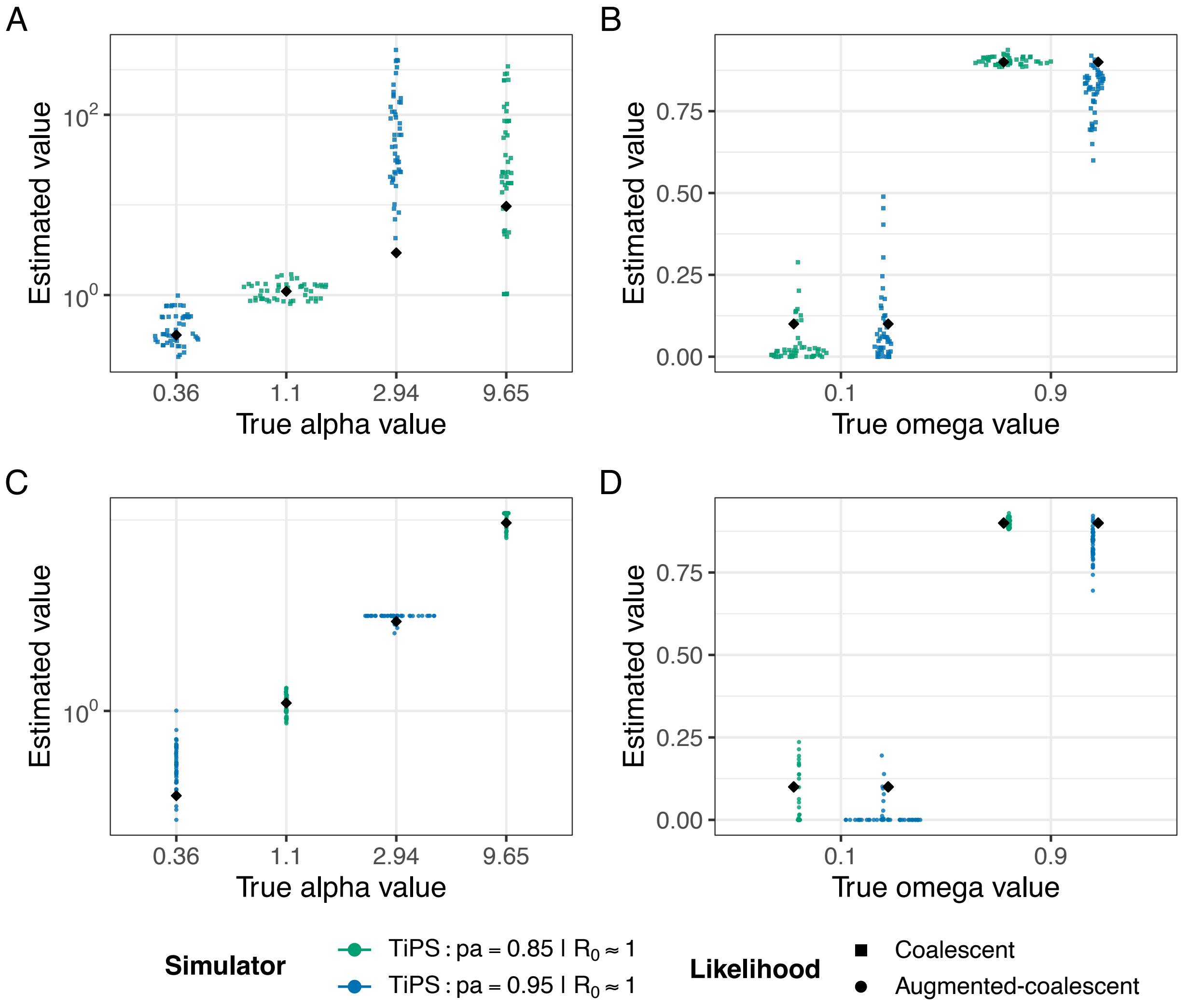


Figure S8: **Sina plot for *TiPS* simulations with 500 tips**: True versus estimated (A) $\alpha$ and (B) $\omega$ values using the (A) and (B) coalescent and (C) and (D) augmented-coalescent likelihoods. Black diamond shapes in the plot are the true values. Depending on combination of parameter values, the proportion of individuals carrying the ancestral state ($p_a$) could be 0.85 or 0.95. For these simulations, we sampled at equilibrium and $R_0$ is expected to be close to 1.0.

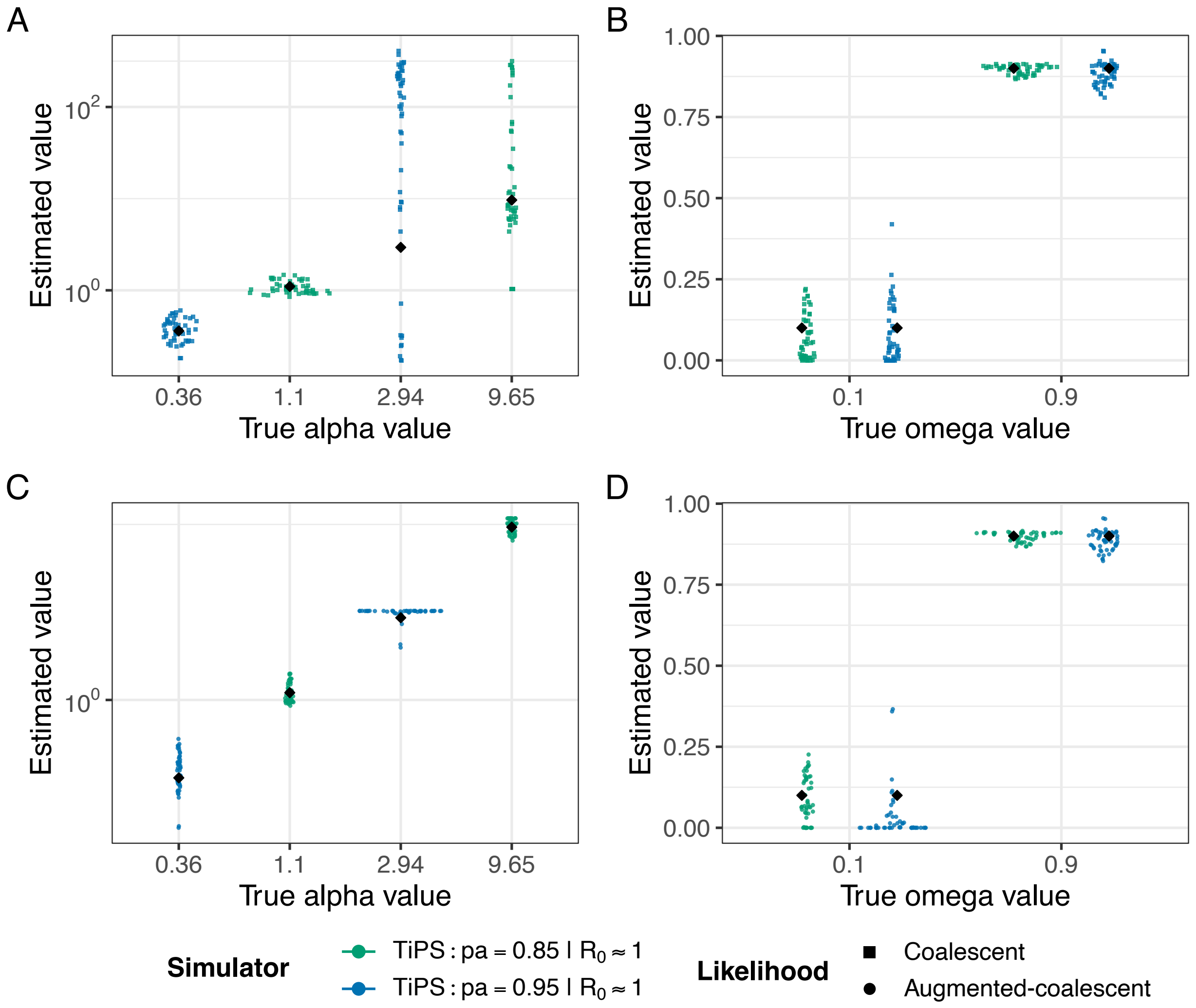


Figure S9: **Sina plot for *TiPS* simulations with 1000 tips**: True versus estimated (A) $\alpha$ and (B) $\omega$ values using the (A) and (B) coalescent and (C) and (D) augmented-coalescent likelihoods. Black diamond shapes in the plot are the true values. Depending on combination of parameter values, the proportion of individuals carrying the ancestral state ($p_a$) could be 0.85 or 0.95. For these simulations, we sampled at equilibrium and $R_0$ is expected to be close to 1.0.

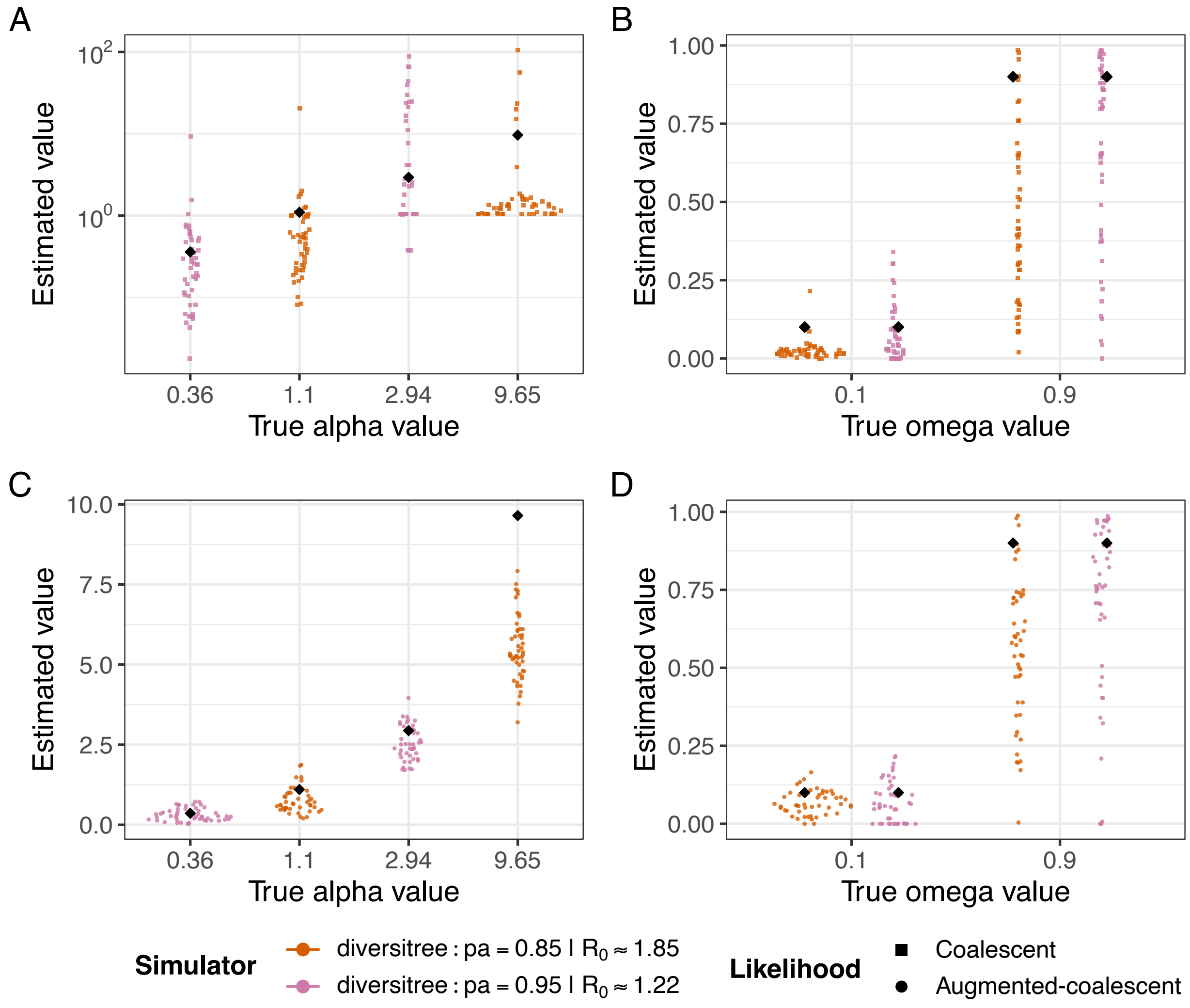


Figure S10: **Sina plot for *diversitree* simulations with 500 tips**: True versus estimated (A) $\alpha$ and (B) $\omega$ values using the (A) and (B) coalescent and (C) and (D) augmented-coalescent likelihoods. Black diamond shapes in the plot are the true values. Depending on combination of parameter values, the proportion of individuals carrying the ancestral state ($p_a$) could be 0.85 or 0.95. We grouped the values of $R_0$ into two groups by averaging the observed $R_0$ to 1.22 and 1.85 (see Table S2).

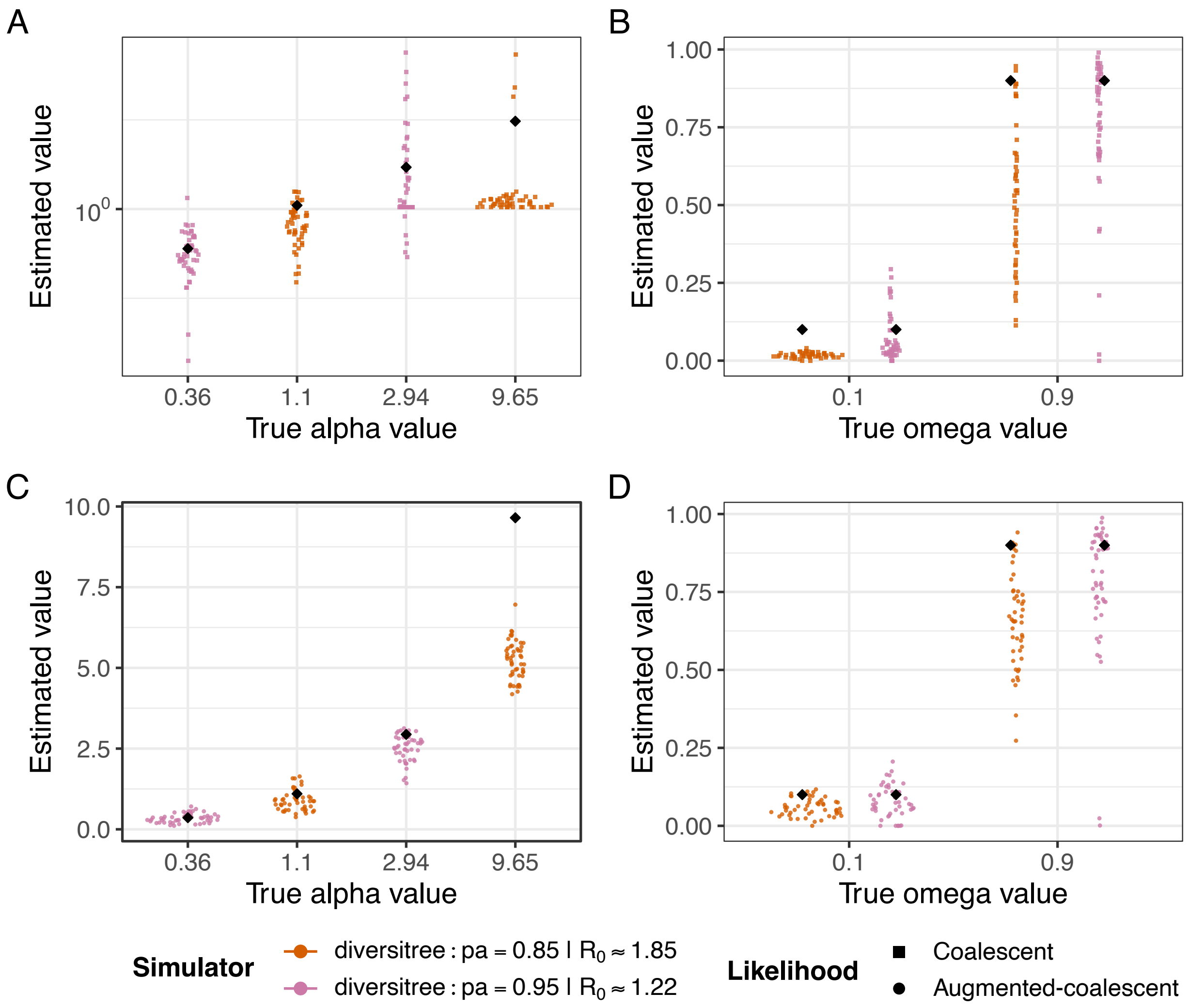


Figure S11: **Sina plot for *diversitree* simulations with 1000 tips**: True versus estimated (A) $\alpha$ and (B) $\omega$ values using the (A) and (B) coalescent and (C) and (D) augmented-coalescent likelihoods. Black diamond shapes in the plot are the true values. Depending on combination of parameter values, the proportion of individuals carrying the ancestral state ($p_a$) could be 0.85 or 0.95. We grouped the values of $R_0$ into two groups by averaging the observed $R_0$ to 1.22 and 1.85 (see Table S2).

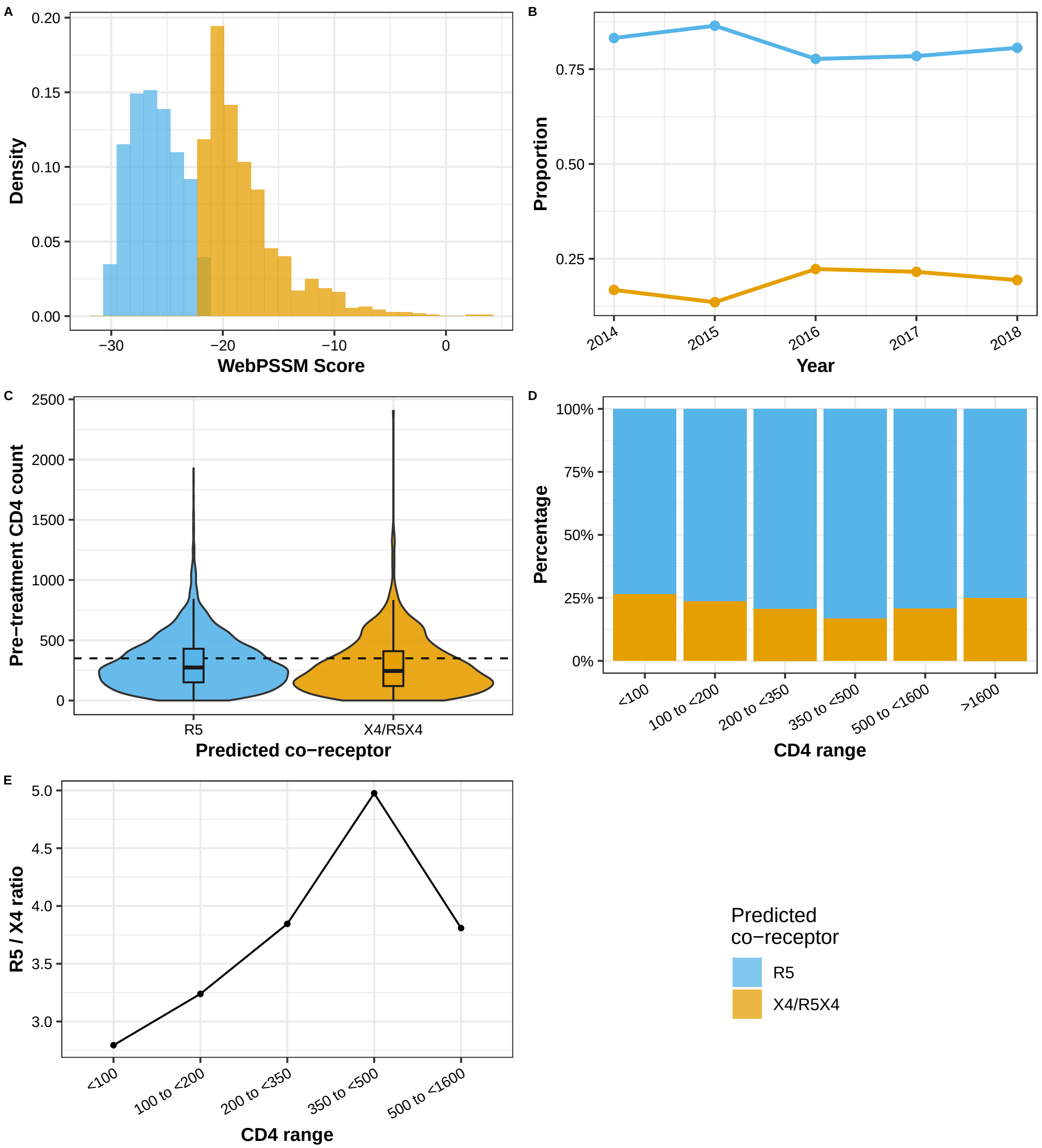


Figure S12: **Co-receptor tropism predictions and association with CD4 counts for the PopART Zambian cohort.** (A) Distribution of WebPSSM scores across predicted R5 and X4/R5X4 viruses. (B) Trend in co-receptor usage over time. (C) Violin and boxplot of individual-level pre-treatment CD4 count measured at visit for the two predicted co-receptors, with the clinically relevant threshold of 350 cells/mm$^3$ represented by the horizontal line. (D) Percentage of individuals infected by R5 and X4/R5X4 viruses for different CD4 ranges. (E) Ratio of individuals infected by R5 vs X4/R5X4 for different CD4 ranges.

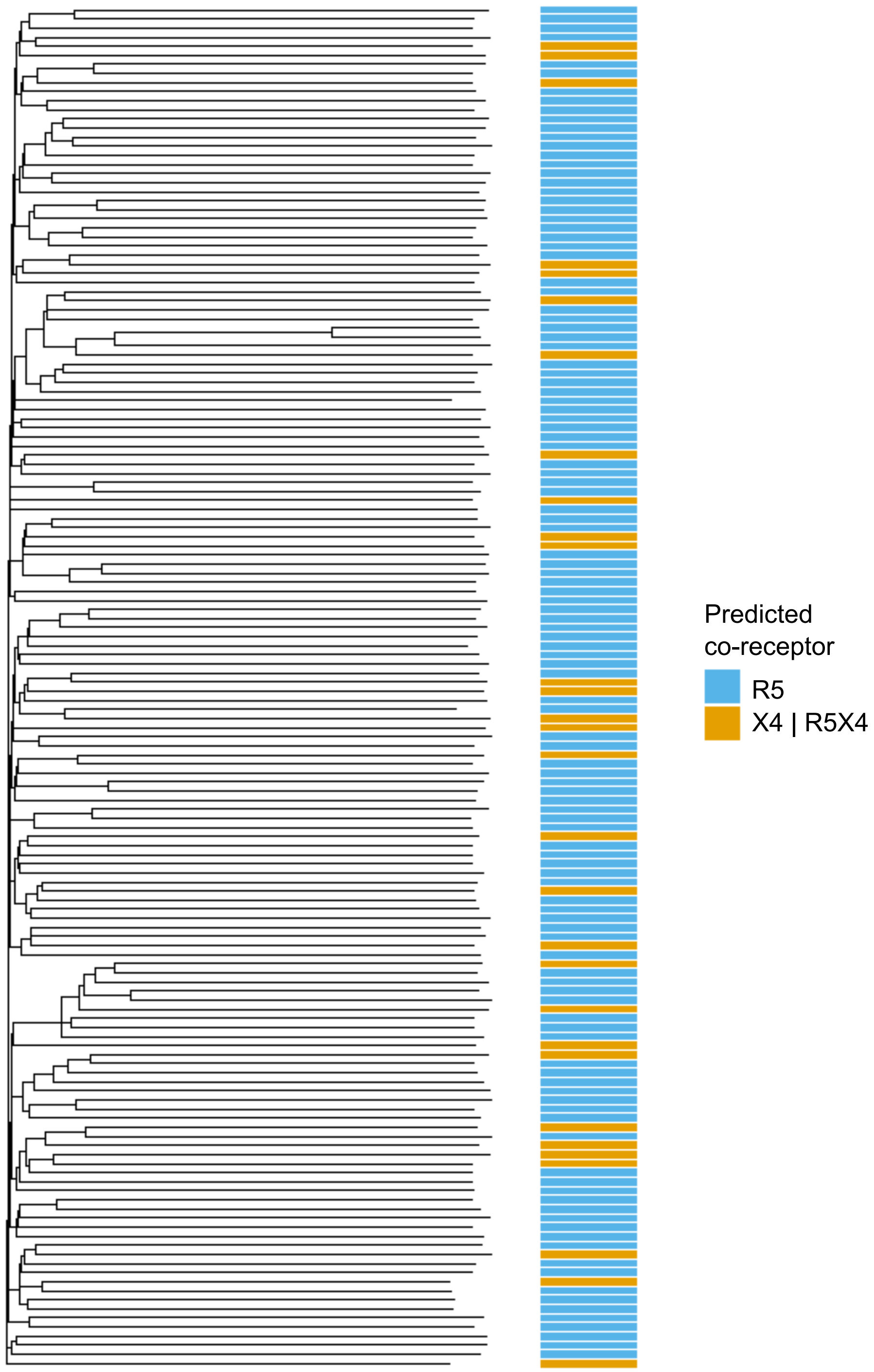


Figure S13: **Phylogeny for HIV *env* (with variable loops (V1-V5) masked) for PopART Zambian cohort.** The phylogeny includes 150 tips subsampled from the larger 4056 tip phylogeny analysed in this study. The predicted R5 and X4|R5X4 viruses are indicated and show X4 or dual-tropic occurrences separately across the phylogeny as well as some clustering. This results from the relative fitness of the two phenotypes at the within-host level ($\alpha \approx 6.9$) and between-host level ($\omega \approx 0.1$), as discussed in the Results section in the main text.
